# Structural Brain Co-Maturation in the Neonatal Period

**DOI:** 10.64898/2026.09.20.752950

**Authors:** Clara F Weber, Emma C Robinson, Logan ZJ Williams, Sofie L Valk

**Affiliations:** Max Planck Institute for Human Cognitive and Brain Sciences, Leipzig, Germany; Institute for Neurosciences and Medicine-7, Forschungszentrum Jülich, Jülich, Germany; Research Department of Biomedical Computing, School of Biomedical Engineering and Imaging Sciences, King’s College, London, United Kingdom; Heinrich-Heine-Universität, Düsseldorf, Germany

## Abstract

**Background:** The neonatal period is characterized by neurodevelopmental processes coinciding with rapid physiological transitions, however, spatial patterns of brain expansion in early life remain incompletely characterized. Here, we leverage grey matter structural covariance in term- and preterm-born neonates to study the interregional synchrony of neonatal brain maturation along principal brain organizational axes.

**Methods:** We used structural MRI data from the developing Human Connectome Project (n=525; 435 term-born) scanned at 37-44 weeks postmenstrual age (PMA)), stratifying the sample by term status and PMA. Based on cortical thickness (CT) and surface area (SA) metrics, we computed age group-(CV_g_) and individual-level (CV_i_) structural covariance matrices. Applying surface-based linear models, we identified changes in interregional covariance with postmenstrual age (PMA) and gestational age at birth (GAB), and contextualized alterations to neonatal functional networks. Using gradient decomposition, we identified major trajectories of cortical expansion on individual and group levels, and determined their correlation to archetypal geometric, evolutionary, and functional axes.

**Results:** We found age-related changes reflecting cortical expansion and an increased structural differentiation with age. Gradient decomposition of group-level covariance revealed that primary components followed both an posterior-anterior distribution, anchored in the visual cortex, and a sensorimotor-association gradient. Individual-level covariance gradients adhered most to geometric axes, and diverged from a sensorimotor-association pattern with increasing PMA. Term-born neonates exhibited a higher spatial correlation to functional and geometric axes than preterm-born neonates, overall pointing to different expansion patterns in neonates of the same PMA.

**Conclusion:** We found trajectories of structural maturation in the neonatal period that appear coordinated along geometric and functional axes. On an age group level, term-born neonates appear to follow archetypal axes more than preterm-borns at term-equivalent age, underlining differential expansion patterns in preterm-born neonates.

## INTRODUCTION

Neurodevelopment is a complex and dynamic process unfolding across embryogenesis, childhood, and adolescence [1], [2]. While basic brain structures and functions develop in utero, the brain remains immature at birth [3], [4]. This altriciality characterizes human development and potentially facilitates extended malleability of neural circuits [5], [6]. At the same time, the developing brain’s immaturity renders it especially susceptible to external influences [7], [8], [9]. The perinatal period, encompassing late fetal and early neonatal stages, represents a time of rapid neurobiological change [10], [11], [12]. The transition from an intrauterine to an extrauterine environment induces a cascade of physiological shifts towards metabolic and respiratory autonomy, alongside circulatory pressure changes and autonomous thermoregulation [12], with concurrent exposure to rich multisensory stimulation and novel social and microbial environments [13]. These changes coincide with critical neurodevelopmental processes such as neuronal and glial proliferation, migration, synaptogenesis, and myelination [14], [15], [16], [17].

Pregnancies vary in length, and neonates are thus exposed to birth-related changes at different stages of brain maturity [14], [18]. Gestational age at birth (GAB) quantifies the timespan of intrauterine development and helps to describe the neurodevelopmental stage at birth. Clinically, GAB is used to determine maturity: neonates born at or before 36+6w gestation are considered *preterm*, whereas a typical *term* pregnancy spans 37-41 gestational weeks [18]. Postmenstrual age (PMA), i.e., age from onset of gestation, aggregates GAB and chronological age, thus representing the total developmental timespan. Within neonatal cohorts matched for PMA, infants have spent different time spans ex utero (chronological age) depending on their GAB [3], [19]. For example, two infants, 14 and four weeks old but born at 28 and 38 weeks, respectively, share the same PMA of 42 weeks (Infant 1: 28w GAB + 14w chronological age = 42w PMA, infant 2: 38w GAB + 4w chronological age = 42w PMA; **Fig. 1A**). Thus, while both brains underwent the same overall length of brain development, different percentages of this time were spent in intra- and extrauterine environments, and they underwent physiological changes and transition to extrauterine stimuli at different developmental stages [8], [14]. Birth timing can alter neurodevelopmental trajectories: Premature birth has been associated with white matter and myelination deficits manifesting in atypical connectivity patterns [4], [9], [10], [20].

**Figure 1.**
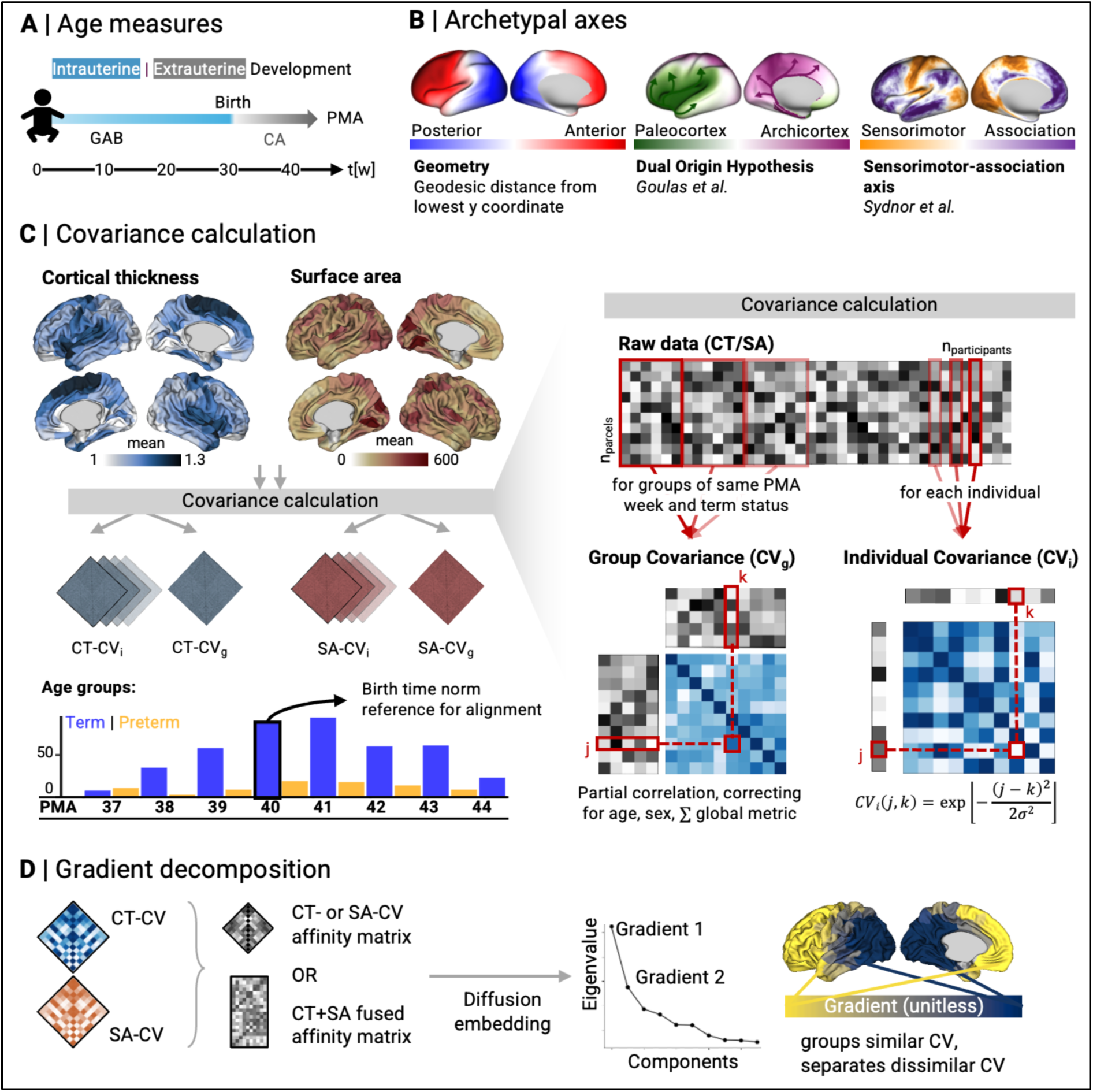
**A** Age metrics for an example neonate. Intra- and extrauterine development are shown in blue and grey, respectively. **B** Archetypal axes of development. Left: posterior-anterior geometric constraints, middle: Dual Origin hypothesis, pattern reproduced from Goulas et al., right: sensorimotor-association axis by Sydnor et al. **C** Mean CT and SA metrics across all groups; flowchart, computation of group-level and individual covariance from CT and SA metrics: CVg and CVi were computed for CT and SA separately. Figure shows how CV was determined based on values in parcels j, k. **D** Analysis workflow: Matrix fusion and dimensionality reductions for gradient decomposition. CT = Cortical thickness, CT-CVg = group-level covariance calculated from cortical thickness, CT-CVi = individual covariance calculated from cortical thickness, GAB = Gestational age at birth, ICV = intracranial volume, PMA = postmenstrual age, SA = surface area, SA-CVg = group-level covariance calculated from surface area, SA-CVi = individual covariance calculated from surface area.

Continued neurodevelopment builds the basis for integrated functional processing across cortical hierarchies [16], [21], supported by subtle variations in cortical structure. These organized structural axes are refined throughout development by intrinsic genetic programming [22], activity dependent plasticity and biology-environment interactions [14], [23], [24].The perinatal period is characterized by a rapid incline in synaptogenesis, followed by myelination and volume expansion [16], [17]. Throughout later childhood and adolescence, the abundant connections formed during embryogenesis and infancy undergo pruning and differentiation, converging into an increasingly efficient network of scarce but meaningful neuronal connections [25]. The resulting spatial variations in connectivity patterns and microstructural differentiation enable functional segregation and integration of information processing, thus enhancing computational power and efficiency [26], [27].

Across neurodevelopment, brain maturation physiologically follows major spatiotemporal axes. A primary spatial pattern of brain function unfolds that juxtaposes sensorimotor areas, handling information in-/output, and brain regions governing higher-order cognition, such as self-generated thought, on different ends of an archetypal sensorimotor-association-axis [1], [28], [29], [30]. Topographically, this axis runs from visual and sensorimotor areas towards prefrontal and temporal regions, and the temporoparietal junction (**Fig. 1B)** [1], [29]. The sensory-association axis may be additionally refined by allo-cortical patterning centers, as described by the Dual Origin theory [23], [31], [32], [33], [34]. and expanded by the multinodal induction–exclusion in network development model [35]. Here brain developmental patterns are thought to originate not from sensory patterning centers alone but also stem from the hippocampus (archicortical origin) and piriform cortex (paleocortical origin) and radiate outwards, resulting in an posterior-anterior and medial-lateral distribution of developmental columns (**Fig. 1B)** [34], [35]. Finally, development also is organized along an posterior-anterior, or rostro-caudal, axis, linked to neurogenesis [36]. Understanding how these archetypal large-scale trajectories restrain and guide neonatal brain expansion can help identify early alterations and give insights into a foundational period of brain development.

Covariance, i.e., markers of similarity between brain metrics within an individual or a group, has been used to describe which regions are most similar or different from one another [23], [37], [38]. Brain regions of similar genetic and microarchitectural composition show high covariance with one another; likewise, high covariance may indicate shared function, as the laminar and cellular setup of a brain region is closely tied to its localization along functional brain trajectories [22], [43], [44], [45]. As such, the covariance of brain morphology markers discloses how brain regions differentiate and provides a proxy for the brain’s increasing spatial differentiation across development [40], [42], [43]. Covariance can be assessed as partial correlation of brain metrics across a population, showing large-scale trends of regional co-maturation [23]. Similarly, individual-level covariance, based on a similarity measure between every pair of parcels and normalized by overall variance [38], can reflect the differentiation of cortical measures on an individual scale, allowing for the contextualization of covariance patterns with subject-specific granularity and the consideration of environmental mediators. Both metrics were first introduced based on cortical thickness (CT) measurements [23], [44]. In neonates, CT is less robust than in adults due to the immaturity of tissues and obtuse grey-white matter boundary [45], [46]; thus, complementation with surface area (SA) metrics potentially contributes to determining robust trajectories of structural co-maturation in this age group, offering insights from easily obtainable imaging markers compared to microstructural profiling approaches requiring specialized imaging acquisition [47], [48].

Here, we aim to investigate how birth timing shapes coordinated cortical development along major brain organizational axes. To this end, we examine the covariance of brain morphological markers derived from structural imaging, CT and SA, to construct age-specific representations of brain architecture. By comparing these patterns to normative configurations of cortical maturation, we seek to provide a detailed description of structural brain maturation in early neurodevelopment, thereby improving our understanding of early-life determinants of brain architecture. In addition, we contextualized early neurodevelopment to frameworks of hierarchical brain organization and aim to disentangle growth effects along sensory-association, allocortical, and posterior-anterior dimensions.

## METHODS

### Participants

We leveraged data from the developing Human Connectome Project (dHCP), an open-data imaging initiative led by King’s College London, Imperial College London, and Oxford University [49]. We used data from the third release, corresponding to an acquisition window of 2014-2019. Gestational age was determined from the last menstrual period and confirmed by prenatal ultrasound, where applicable. Exclusion criteria for enrollment in dHCP encompassed scanning contra-indications, e.g., due to implanted devices, intolerance of the scanning period despite intensive care, and parent language barriers that did not allow for informed consent to participation. We considered those scans acquired beginning at term-equivalent age (37-44w PMA). Subjects with reports of congenital abnormalities and twin-twin-transfusion syndrome were excluded. To delineate developmental effects, we aggregated subjects by their PMA, resulting in eight age groups containing data from infants scanned at 37-44 weeks (**Table 1**). Of note, no data were available for preterm infants scanned at 44 weeks PMA after applying exclusion criteria.

**Table 1.** Baseline demographic information for all participants in PMA bins. Apgar scores were not available for all subjects. GAB = gestational age at birth, PMA = postmenstrual age, SD = standard deviation

| Week PMA | all | 37 | 38 | 39 | 40 | 41 | 42 | 43 | 44 |
| --- | --- | --- | --- | --- | --- | --- | --- | --- | --- |
| **n (all)** | 525 | 21 | 40 | 69 | 109 | 114 | 76 | 72 | 24 |
| Female, n (%) | 233 (44%) | 11 (52%) | 10 (25%) | 24 (35%) | 47 (43%) | 56 (49%) | 34 (45%) | 34 (47%) | 17 (71%) |
| PMA, mean±SD | 41.13±1.73 | 37.52±0.33 | 38.47±0.28 | 39.45±0.26 | 40.5±0.29 | 41.43±0.28 | 42.38±0.33 | 43.42±0.28 | 44.24±0.25 |
| GAB, mean±SD | 38.55±3.51 | 36.77±0.77 | 37.36±2.37 | 38.26±2.43 | 38.4±3.59 | 38.78±4.06 | 38.61±4.05 | 39.12±3.89 | 40.57±0.96 |
| **1min Apgar** |  |  |  |  |  |  |  |  |  |
| 7-10, n (%) | 415 (81%) | 19 (90%) | 35 (90%) | 59 (86%) | 90 (83%) | 85 (77%) | 58 (77%) | 57 (83%) | 12 (67%) |
| 4-6, n (%) | 60 (12%) | 2 (10%) | 4 (10%) | 8 (12%) | 13 (12%) | 16 (14%) | 11 (15%) | 4 (6%) | 2 (11%) |
| 0-3, n (%) | 36 (7%) | 0 (0%) | 0 (0%) | 2 (3%) | 6 (6%) | 10 (9%) | 6 (8%) | 8 (12%) | 4 (22%) |
| **5min Apgar** |  |  |  |  |  |  |  |  |  |
| 7-10, n (%) | 478 (94%) | 21 (100%) | 38 (97%) | 67 (97%) | 105 (96%) | 104 (94%) | 66 (88%) | 63 (91%) | 14 (78%) |
| 4-6, n (%) | 10 (2%) | 0 (0%) | 0 (0%) | 1 (1%) | 2 (2%) | 1 (1%) | 3 (4%) | 2 (3%) | 1 (6%) |
| 0-3, n (%) | 23 (5%) | 0 (0%) | 1 (3%) | 1 (1%) | 2 (2%) | 6 (5%) | 6 (8%) | 4 (6%) | 3 (17%) |
| **n (term)** | 435 | 9 | 36 | 59 | 89 | 95 | 61 | 62 | 24 |
| Female, n (%) | 193 (44%) | 5 (56%) | 10 (28%) | 22 (37%) | 36 (40%) | 47 (49%) | 25 (41%) | 31 (50%) | 17 (71%) |
| PMA, mean±SD | 41.22±4.77 | 37.74±0.1 | 38.47±0.28 | 39.43±0.26 | 40.54±0.27 | 41.43±0.28 | 42.35±0.31 | 43.43±0.28 | 44.24±0.25 |
| GAB, mean±SD | 39.90±1.19 | 37.29±0.15 | 38.08±0.47 | 39.12±0.4 | 39.92±0.8 | 40.42±1.01 | 40.38±1.06 | 40.54±1.06 | 40.57±0.96 |
| **1min Apgar** |  |  |  |  |  |  |  |  |  |
| 7-10, n (%) | 360 (85%) | 8 (89%) | 32 (89%) | 54 (92%) | 78 (88%) | 73 (79%) | 50 (83%) | 53 (88%) | 12 (67%) |
| 4-6, n (%) | 36 (9%) | 1 (11%) | 4 (11%) | 5 (8%) | 6 (7%) | 11 (12%) | 5 (8%) | 2 (3%) | 2 (11%) |
| 0-3, n (%) | 27 (6%) | 0 (0%) | 0 (0%) | 0 (0%) | 5 (6%) | 8 (9%) | 5 (8%) | 5 (8%) | 4 (22%) |
| **5min Apgar** |  |  |  |  |  |  |  |  |  |
| 7-10, n (%) | 400 (95%) | 9 (100%) | 35 (97%) | 58 (98%) | 87 (98%) | 87 (95%) | 54 (90%) | 56 (93%) | 14 (78%) |
| 4-6, n (%) | 4 (1%) | 0 (0%) | 0 (0%) | 1 (2%) | 1 (1%) | 0 (0%) | 1 (2%) | 0 (0%) | 1 (6%) |
| 0-3, n (%) | 19 (4%) | 0 (0%) | 1 (3%) | 0 (0%) | 1 (1%) | 5 (5%) | 5 (8%) | 4 (7%) | 3 (17%) |
| **n (preterm)** | 90 | 12 | 4 | 10 | 20 | 19 | 15 | 10 | 0 |
| Female, n (%) | 41 (46%) | 6 (50%) | 0 (0%) | 2 (20%) | 11 (55%) | 9 (47%) | 9 (60%) | 3 (30%) | - |
| PMA, mean±SD | 40.69±1.85 | 37.35±0.34 | 38.5±0.25 | 39.56±0.24 | 40.33±0.32 | 41.42±0.28 | 42.5±0.4 | 43.33±0.3 | - |
| GAB, mean±SD | 32.00±3.67 | 36.38±0.83 | 30.89±2.91 | 33.2±3.23 | 31.64±3.36 | 30.55±3.53 | 31.41±3.7 | 30.33±3.5 | - |
| **1min Apgar** |  |  |  |  |  |  |  |  |  |
| 7-10, n (%) | 55 (63%) | 11 (92%) | 3 (100%) | 5 (50%) | 12 (60%) | 12 (63%) | 8 (53%) | 4 (44%) | - |
| 4-6, n (%) | 24 (27%) | 1 (8%) | 0 (0%) | 3 (30%) | 7 (35%) | 5 (26%) | 6 (40%) | 2 (22%) | - |
| 0-3, n (%) | 9 (10%) | 0 (0%) | 0 (0%) | 2 (20%) | 1 (5%) | 2 (11%) | 1 (7%) | 3 (33%) | - |
| **5min Apgar** |  |  |  |  |  |  |  |  |  |
| 7-10, n (%) | 78 (89%) | 12 (100%) | 3 (100%) | 9 (90%) | 18 (90%) | 17 (89%) | 12 (80%) | 7 (78%) | - |
| 4-6, n (%) | 6 (7%) | 0 (0%) | 0 (0%) | 0 (0%) | 1 (5%) | 1 (5%) | 2 (13%) | 2 (22%) | - |
| 0-3, n (%) | 4 (5%) | 0 (0%) | 0 (0%) | 1 (10%) | 1 (5%) | 1 (5%) | 1 (7%) | 0 (0%) | - |

**Table 2.**
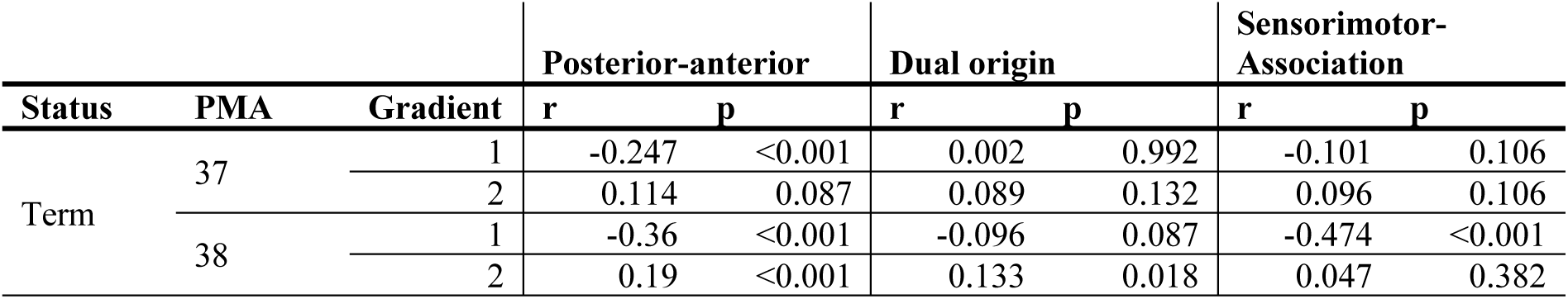
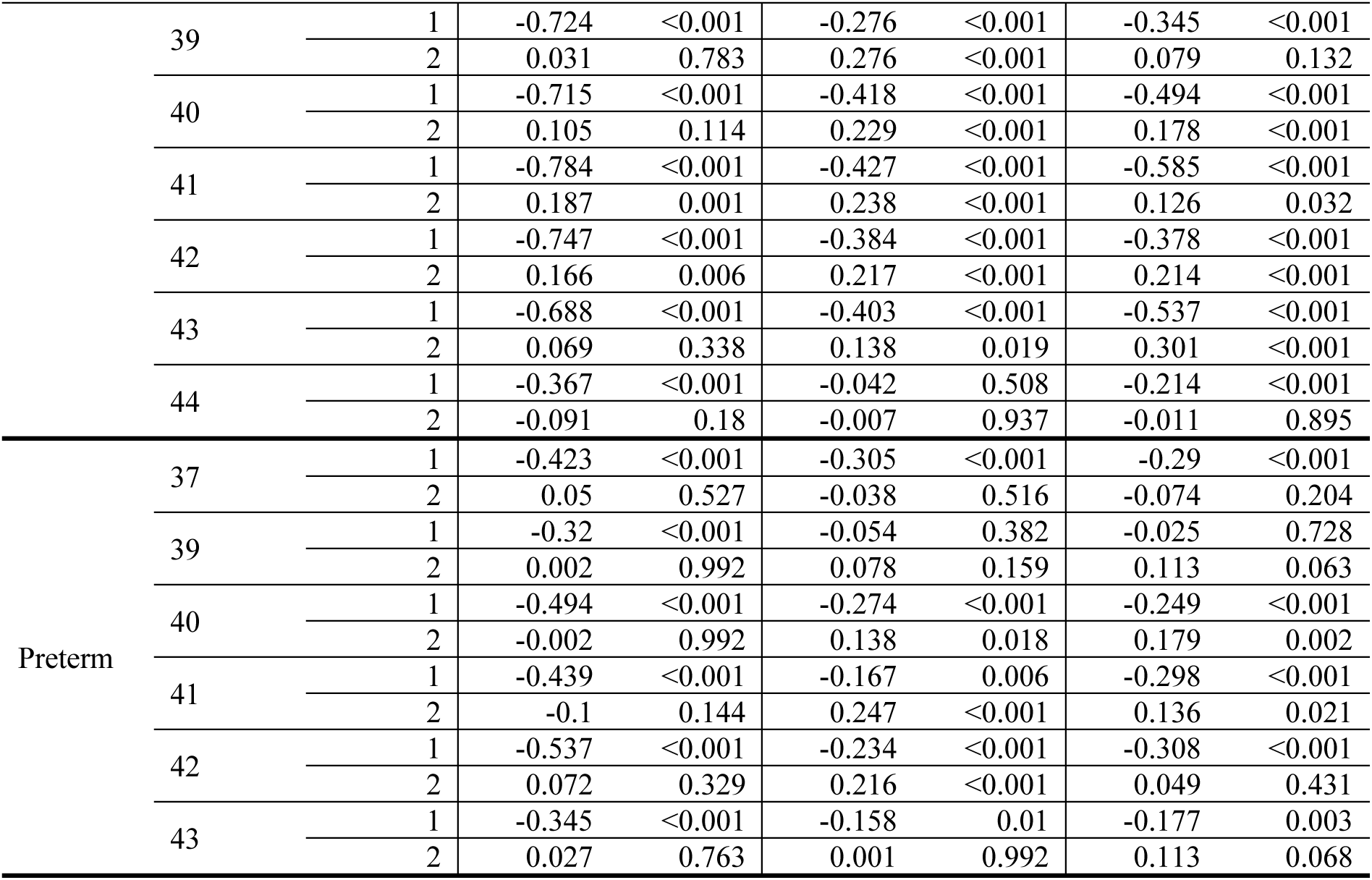
Spearman correlation, tested for spatial autocorrelation in 5,000 variogram-based permutations and adjusted for multiple testing using false discovery rate correction, between age-specific CVg gradients and archetypal axes, corresponding to Figure 3D. Preterm-born neonates in the PMA groups of 38 and 44 weeks were excluded due to insufficient sample sizes.

### Image acquisition

Structural imaging data were acquired using a 3 Tesla Philips Achieva system in a single scan session at the Evelina Newborn Imaging Center of Evelina London Children’s Hospital. Participating neonates were fed and swaddled prior to scanning; no sedation was used, and vital parameters were monitored by a supervising nurse or physician. T2w was acquired with a repetition time of 12 s, echo time of 156 ms, resolution 0.8 x 0.8 mm at 1.7 mm slice thickness, as previously described in detail [50].

### Data processing

Data was processed using the dHCP minimal processing pipeline, previously described by Makropoulos et al. [19]. Briefly, images were bias corrected, and non-brain tissue volumes were discarded. Grey-white matter outlines were delineated and extended outwards to the grey matter-cerebrospinal fluid boundary. Surfaces were then inflated and projected to a spherical body for coregistration. CT was computed as the Euclidean distance between the pial and white matter surfaces, and SA was derived from the pial surface within each parcel. As previously implemented in the HCP minimal pre-processing pipeline, curvature was regressed out from CT and SA to correct for folding bias [51]. All surface-level data was mapped to 300 cortical parcels (150 per hemisphere) [52].

### Covariance calculation

Based on cortical thickness (CT) and surface area (SA) estimates, we calculated two types of structural covariance (**Fig. 1C**): (i) *Group-level covariance* (CV_g_), defined as the partial correlation of group-level metrics in a parcel, while correcting for sex, PMA and global CT or SA values within each participant [23]; (ii) *Individual covariance* (CV_i_), a similarity metric defined by Wee et al. [38] as 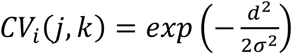 where *CV_i_* (*j*, *k*) represents CV_i_ between parcels *j* and *k*; *d* represents the absolute difference between metrics in parcels *j, k*; and 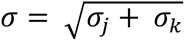, where *σ*_j,*k*_ represents the group-level standard deviation of metrics within parcels *j, k* [38]. Briefly, this method transforms the absolute difference between two parcels to a dissimilarity measure between 0 and 1, where similarity decreases exponentially with absolute difference. As such, high CV_i_ indicates high similarity between two parcels, and decreases with growing dissimilarity. Both CV metrics were determined for SA and CT as individual metrics and are henceforth abbreviated as CT-CV_i/g_ and SA-CV_i/g_, respectively (**Fig 1C**).

### Statistical analysis

On a group level, we fit a linear model *Parcel* = *β*_0_ + *β*_1_ ∗ *PMA* + *β*_2_ ∗ *GAB* + *β*_3_ ∗ *Sex* + *β*_4_ ∗ *global measure* to each surface parcel for each metric available at individual-level granularity (CT, CT-CV_i_, SA, SA-CV_i_) separately. *Global measure* denotes the mean of the respective metric across all cortical parcels. We then determined the effects of PMA and GAB, controlling for multiple comparisons using false discovery rate correction. Parcels with significant changes were then contextualized to neonatal functional networks, considering the intrinsic functional networks described by Molloy and Saygin [53] (**Fig. 2B**). In addition to age, we tested the influence of Apgar scores documented at one and five minutes on brain structural metrics. Briefly, Apgar scores are clinical scores reflecting the overall adaptation of a newborn to the extrauterine development, spanning categories of appearance, pulse, irritability, muscle tone, and respiration [54], with higher scores representing a clinically better constitution. Likewise, we determined the influence of delivery method (Forceps/Ventous: instrumental delivery with forceps/suction, CS-Lab: emergency caesarean delivery in labor, CS-noLab: emergency caesarean delivery not in labor, E-CS: elective caesarean section, SemiE-CS: semi-elective caesarean section; as decoded in the dHCP data release) as compared to vaginal delivery. In both Apgar- and delivery mode-specific analysis, we used linear models similar to the age-related approach above.

**Figure 2.**
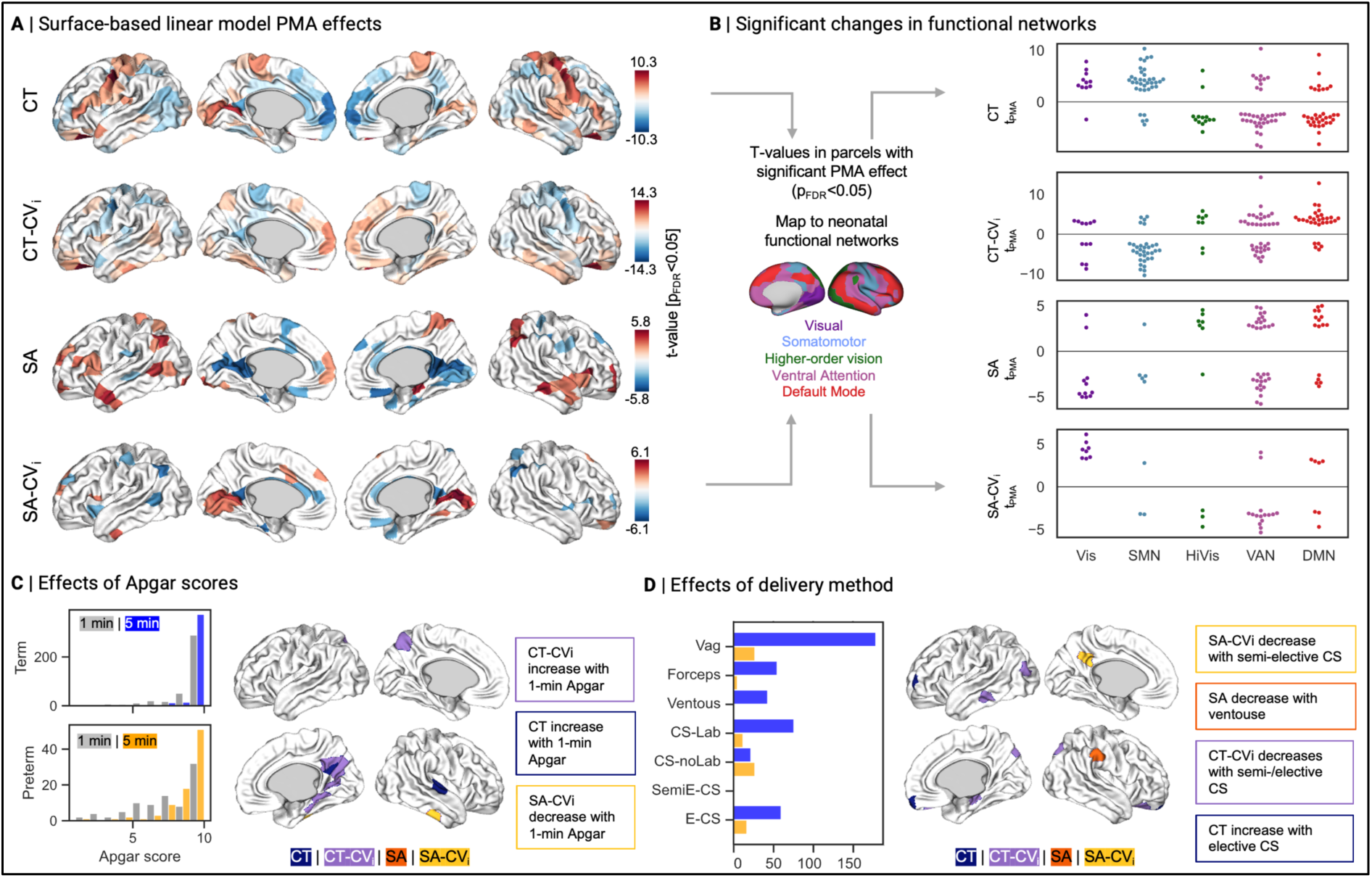
**A** Outcomes of linear model testing for postmenstrual age (PMA) effects while controlling for GAB, sex, and global measure; Brain plots show t-values in statistically significant parcels (p_FDR_<0.05) in four different metrics. **B** Significant parcels plotted by their participation in neonatal functional networks, from left to right Vis = Visual, SMN = Somatomotor network, HiVis = higher-order visual, VAN = ventral attention, DMN = default mode network (Molloy and Saygin). **C** Distribution of Apgar scores at 1 and 5 min in term and preterm neonates, and effects on CT (no significant effects on SA or in 5 min score analysis). **D** Distribution of delivery method (Forceps/Ventous: instrumental delivery with forceps/suction, CS-Lab: emergency caesarean delivery in labor, CS-noLab: emergency caesarean delivery not in labor, E-CS: elective caesarean section, SemiE-CS: semi-elective caesarean section; as decoded in dHCP data release) in term (blue) and preterm (yellow) neonates, and significant effects on CT with elective CS and on SA with ventouse delivery (brain plots show t-values in significant (p_FDR_<0.05) parcels. CT = cortical thickness, CVi = individual covariance, CVg = group covariance, cortical thickness, FDR = false discovery rate, GAB = gestational age at birth, PMA = postmenstrual age, SA = surface area

On an age-group level, i.e., in each PMA week stratum, we tested the group-wise correlation of PMA- and term status-specific CV_g_ to a birth time norm derived from all 40w PMA, term-born subjects. To this end, we determined the row-wise Spearman correlation of each age group’s CV_g_ map to the reference 40w, term-born CV_g_ matrix. We conducted this analysis for both CT-CV_g_ and SA-CV_g_ separately.

### Gradient decomposition

We then decomposed data into gradients, i.e., lower-dimensional axes representing central organizational trajectories in brain structure [55], [56]. **Group level**: For our primary analysis, we fused each PMA group’s CT-CV_g_ and SA-CV_g_ covariance matrices by normalizing and ranking their non-zero values prior to stacking matrices horizontally. The fused gradient in primary analysis will be abbreviated as CT+SA-CV_g_ gradient in the following. Using the *BrainSpace* toolbox [56] (https://github.com/MICA-MNI/BrainSpace), we created an affinity matrix using a normalized angle kernel, and employed diffusion embedding for dimensionality reduction [56]. We then employed Procrustes alignment to a birth time-norm group gradient that was derived from all n=89 term-born, 40 weeks PMA neonates. Of the resulting eight pairs of gradients (term- and preterm-group in each of eight PMA strata), we considered the first and primary gradients as ranked by explained variance for the following analyses. In secondary analyses, we repeated gradient computation for CT-CV_g_ and SA-CV_g_ separately. **Individual level**: Similar to CV_g_ gradients, we then computed CV_i_ gradients at the individual level. To this end, we used a fused CT+SA-CV_i_ matrix in primary analysis, again stacking CT-CV_i_ and SA-CV_i_ matrices prior to diffusion embedding for dimensionality reduction. To make gradients comparable, we derived a reference gradient from a mean covariance matrix of all 40w PMA, term-born infants, and then aligned each individual’s gradient to this template. In parallel to robustness analyses for CV_g_ gradients, we repeated analyses using CT-CV_i_ and SA-CV_i_ separately. To test for potential differences between gradients, we first normalized the unitless local gradient scores by their total range, so that each parcel was assigned its localization within the gradient as a percentage score. In parallel to statistical testing of local CT, SA and respective covariance metrics, we then tested the influence of age on gradient scores. To this end, we employed a surface-based linear model *Parcel* = *β*_0_ + *β*_1_ ∗ *PMA* + *β*_2_ ∗ *GAB* + *β*_3_ ∗ *Sex* to normalized gradient scores, again correcting for multiple comparisons using false discovery rate, to test for any significant changes in brain organizational trajectories related to PMA, GAB and sex. Follow-up analyses considered CT-CV_i_ and SA-CV_i_ gradients in isolation.

### Contextualization to archetypal axes

After gradient decomposition, we contextualized primary and secondary gradients to archetypal brain organizational axes. First, we pre-selected three developmental trajectories that have been shown to guide brain maturation: A geometrically informed posterior-anterior axis, a phylogenetic dual origin model [57], and a functional sensorimotor-association axis [58] (**Fig. 1B**). We constructed reference maps for these three archetypes as follows: For geometry, we first determined normative geodesic distance along the surface of an averaged template 40w surface derived from dHCP [59], [60], [61] using the surface geodesic distance –*all-to-all* command in Connectome Workbench (https://www.humanconnectome.org/software/workbench-command). Geodesic distance, representing proximity along the cortical surface, respects cortical folding patterns and thus reflects the continuity of the cortical surface better than other distance metrics [30]. We determined the mean geodesic distance to the vertex with the lowest y-coordinate in each hemisphere separately, resulting in a posterior-anterior map reflecting geometric constraints. Then, we constructed a phylogenetic map representing dual origins. The dual origin theory postulates cortical growth stemming from archi- and paleocortical origins anchored in the hippocampus and piriform cortex, respectively [34]. To describe this spatial pattern, we projected archi- and paleocortical origins onto the cortical surface, leveraging maps from the PaleoArchiNeo atlas by Zhernovaia et al. [62]. Using normative geodesic distance as described above, we determined the average distance of each cortical vertex to archi- and paleocortical origins, creating a continuous proximity map. Positive values indicate closer proximity to the archicortex; vice versa, negative values show proximity to the paleocortex. Zero-values mark vertices equidistant to either origin (**Fig. 1B**). Given the proposed binary nature of dual origin patterns, we conducted sensitivity analysis in which we utilized a binary reference map, in which every cortical parcel was assigned to the origin it is closest to. For the third archetypal pattern, we leveraged the sensorimotor-association functional pattern from previous work by Sydnor *et al.,* reflective of adult functional cortical organization as a future functional template that neonates are developing towards [1]. Briefly, this map was constructed by aggregating the rank values of ten different cortical brain features (T1w/T2w ratio as a proxy for anatomical hierarchy [63], [64], functional and evolutionary hierarchies [39], [55], [65], allometric scaling [66], cerebral blood flow [67], [68], gene expression [69], cortical thickness [70], externopyramidization [71], and meta-analytic functional maps from NeuroSynth [72], [73]) [1].

After the construction of reference maps, we assessed how gradients at individual-level granularity corresponded to the three pre-selected axes. To this end, we determined the Spearman correlation between each group’s CT+SA-CV_g_ gradient to the posterior-anterior, dual origin, and sensorimotor-association axis, correcting for spatial autocorrelation in 5,000 variogram-based permutations [74], [75]. In robustness analyses, we repeated testing using CT-CV_g_ and SA-CV_g_ gradients separately. We then repeated analysis on the individual level, leveraging each individual’s CT+SA-CV_i_ gradient. Due to the exploratory approach of the analysis on an individual scale, we did not test for statistical significance in these analyses.

### Multimodal normative maps

In addition to the three archetypal axes described above, we explored the correspondence to normative imaging patterns that are collected and made accessible in the *neuromaps* repository [76]. Maps that allowed for transformation into *fslr* surface space and were considered relevant in the context of metabolic, functional, and chemoarchitectural contextualization were selected, with a full list and references given in **Table 3** [67], [68], [77], [78], [79], [80], [81], [82], [83], [84], [85], [86], [87], [88], [89], [90], [91], [92], [93], [94]. We determined the correspondence of individual gradients to microstructural and chemoarchitectural patterns by determining Spearman’s correlation between each individual’s gradient and the respective map. Individual correlation values were then r-to-z transformed, and differential correspondence to trajectories was determined as Cohen’s d effect size measures. Statistical significance between r-to-z transformed correlation coefficients was then assessed using unpaired t-tests, adjusting for multiple comparisons using false discovery rate correction.

**Table 3.**
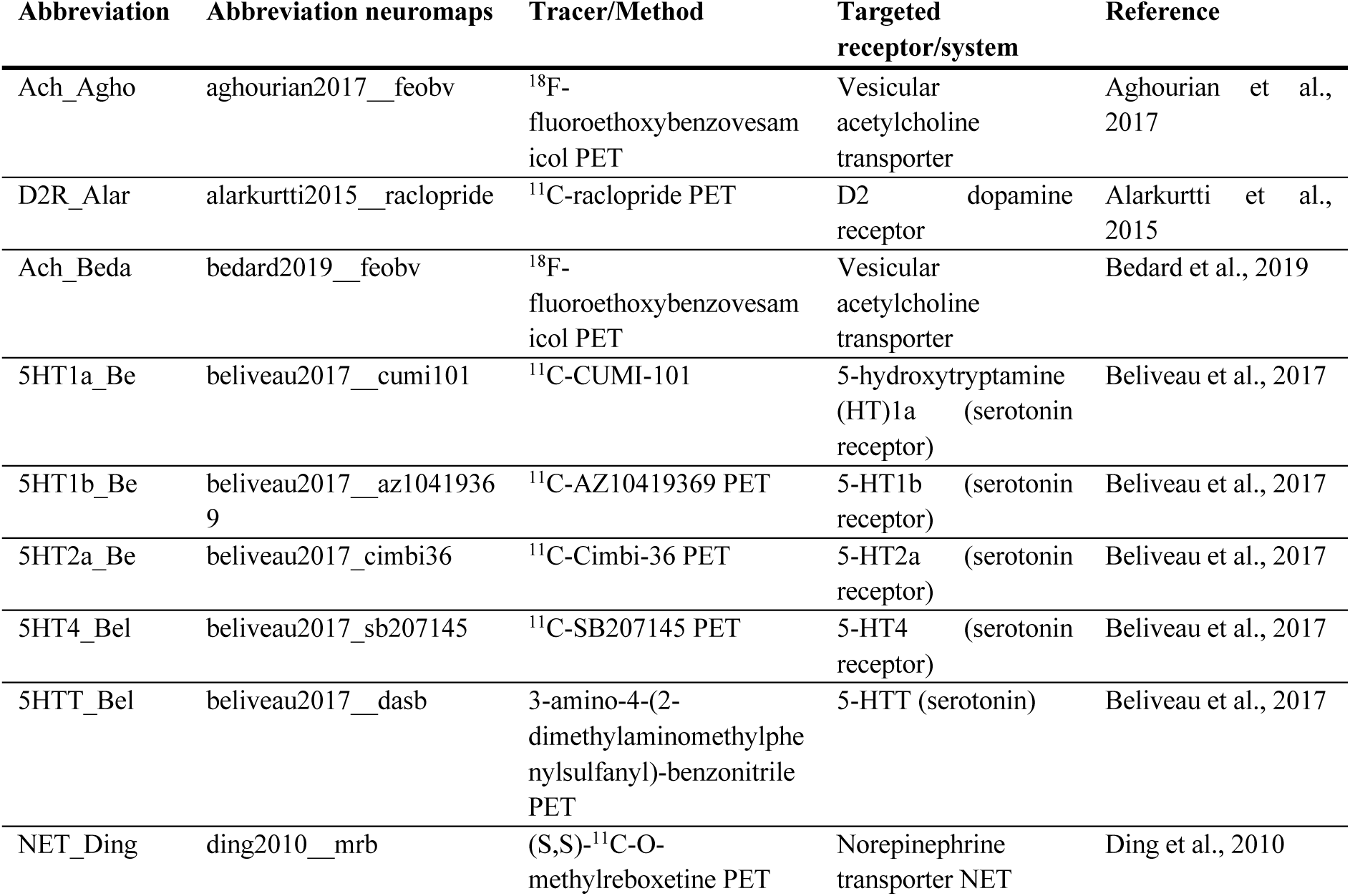
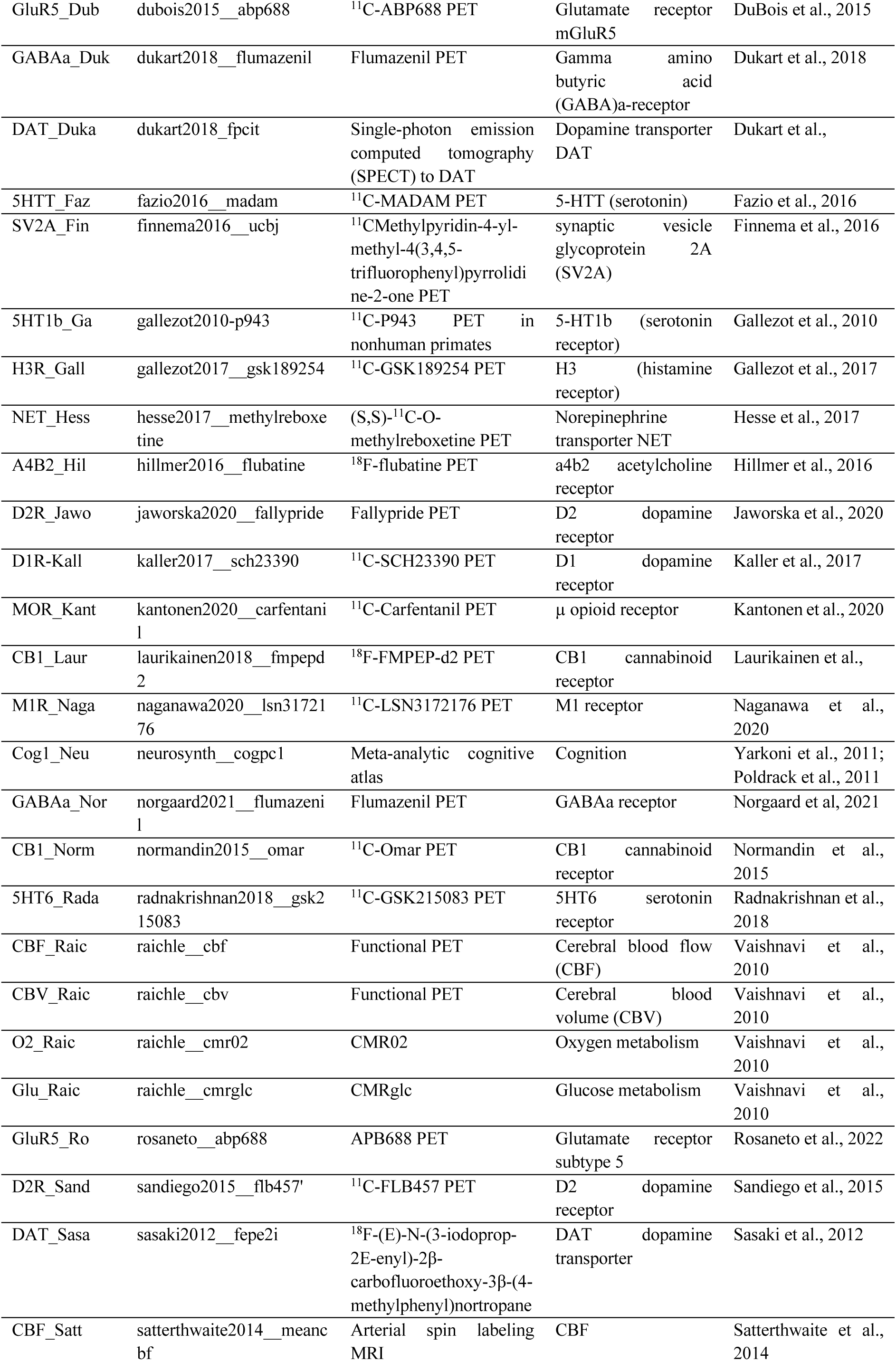
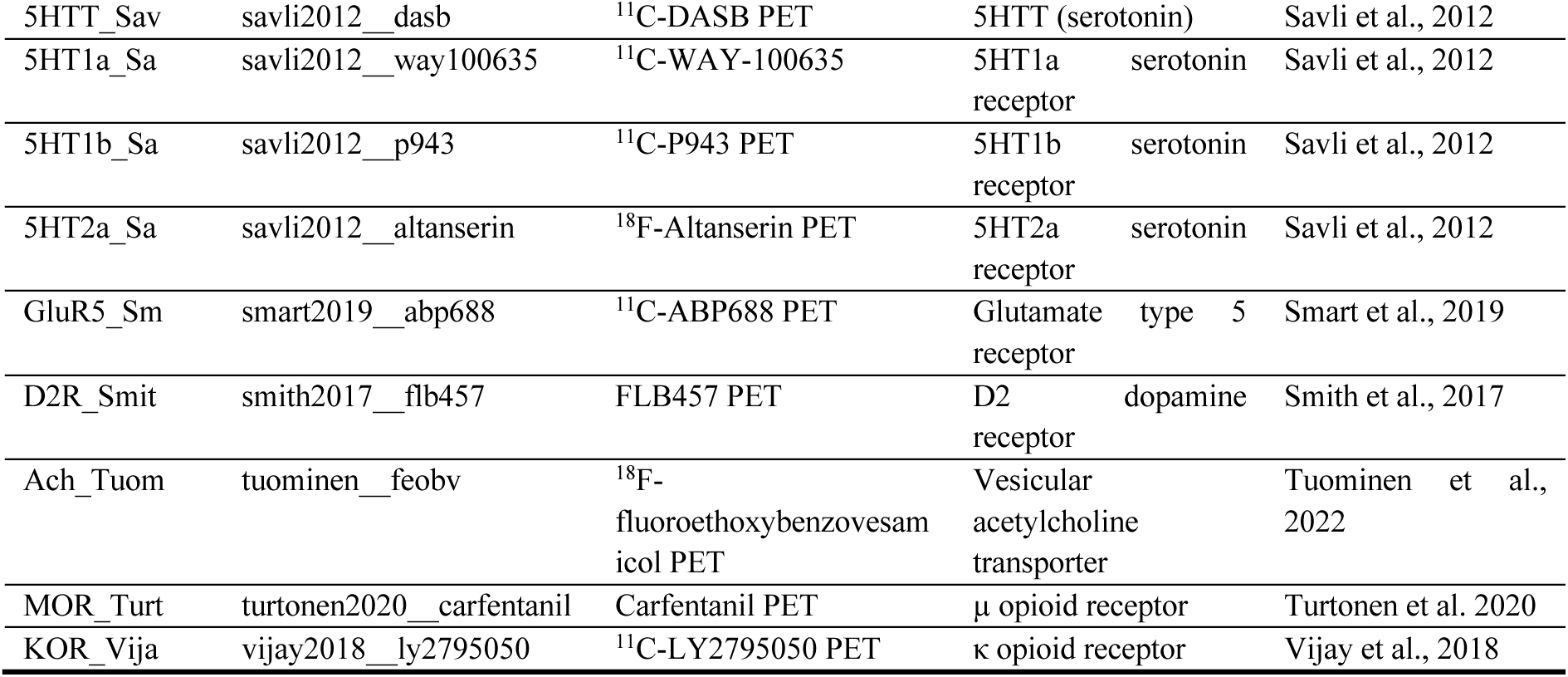
Full list of selected chemoarchitectural maps with references: Aghourian et al., positron emission tomography (PET) tracer binding vesicular acetylcholine transporter using ^18^F-fluoroethoxybenzovesamicol (FEOBV) [77], [94]; Alarkurtti et al., PET tracer binding to D2 dopamine receptor using ^11^C-raclopride [116]; Bedard et al., PET tracer binding to acetycholine transporter using FEOBV [117]; Beliveau et al., PET tracer binding to 5-HT1a, 5-HT1b, 5-HT2a, 5-HT4, and 5-HTT serotonin receptors [92]; Ding et al. PET tracer binding to norepinephrine transporter NET using (S,S)-^11^C-O-methylreboxetine [77], [118], [119], [120], [121]; Dubois et al., PET tracer binding to the glutamate receptor mGluR5 [77], [91]; Dukart et al., PET tracer binding to GABAa-receptor and single-photon emission computed tomography (SPECT) of DAT dopamine transporter [122]; Fazio et al., PET tracer binding to 5-HTT serotonin receptor [90]; Finnema et al., PET tracer binding to synaptic vesicle glycoprotein 2A (SV2A), a synapse marker [89], [123], [124], [125]; Gallezot et al., PET tracer binding to 5-HT1b serotonin receptor and H3 histamine receptor [88]; MRI T1w/T2w ratio myelin map from the Human Connectome Project S1200 release [126]; Hesse et al., PET tracer binding to NET norepinephrine transporter [86]; Hillmer et al., PET tracer binding to a4b2 acetylcholine receptor using ^18^F-flubatine [85]; Jaworska et al., PET tracer binding to D2 dopamine receptor using Fallypride [77], [83]; Kaller et al., PET tracer binding to D1 receptor using ^11^C-SCH23390 [87]; Kantonen et al., PET tracer binding to µ opioid receptors using Carfentanil [127]; Laurikainen et al., PET tracer binding to CB1 cannabinoid receptor [78]; Naganawa et al., PET tracer binding to M1 receptor [128]; Yarkoni et al., first component of cognitive atlas [72], [73]; Norgaard et al., PET tracer binding to GABAa receptors [93]; Normandin et al., PET tracer binding to cannabinoid receptor binding [129]; Vaishnavi et al., functional PET for cerebral blood flow (CBF) and volume (CBV), and oxygen metabolism [68]; Rosaneto et al., PET tracer binding to glutamate receptor type 5 [77]; Sandiego et al., type 2 dopamine receptor binding [82]; Sasaki et al., PET tracer binding to dopamine transporter [81]; Satterthwaite et al., CBF from ASL imaging [67]; Savli et al., multitracer PET binding to serotonin receptors [130]; Smart et al., PET tracer binding to glutamate receptor type 5 [84]; Smith et al., PET tracer binding to dopamine receptor [131]; Tuominen et al., PET tracer binding to acetylcholine transporter [77]; Turtonen et al., PET tracer binding to µ opioid receptor [79] and Vijay et al., PET tracer binding to κ opioid receptor binding [80].

## RESULTS

### Spatial patterns of structural expansion

Across our sample of 525 neonates from dHCP (**Table 1**), CT measurements exhibited the highest values in medial frontal and temporal regions (**Fig. 1C**), and SA showed an posterior-anterior pattern with peaks in the occipital cortex (**Fig. 1C**), in alignment with previous reports [95], [96], [97]. To study temporal effects, we split the sample into age cohorts by PMA, resulting in eight subgroups scanned in weeks 37-44 PMA (**Table 1**). The distribution of CT and SA metrics within these cohorts exhibited a similar spatial distribution, with expanding CT and SA metrics observed with age, resulting in overall increasing values across age. The age cohort-wise distribution is displayed in **supplemental figures 1,2.**

### Age-related changes of structural metrics

Next, we probed how brain regions change and differentiate in the neonatal period. To this end, we calculated group-(CV_g_) and individual-level (CV_i_) covariance metrics for CT and SA metrics, respectively. The workflow for covariance matrix construction is outlined in **Fig. 1C**. The construction of two different covariance metrics allowed to capture complementing effects: CV_g_ captures similarity across brain parcels in a group, thus reflecting overarching patterns of brain maturation that are present on a group level, while CV_i_ captures the similarity of a parcel to other parcels within an individual. Across age cohorts, CT-CV_g_ appeared relatively high in frontal regions, whereas the precuneus and parts of the prefrontal cortex in younger cohorts exhibited higher dissimilarity (**suppl. Fig. 1B)**. Likewise, SA-CV_g_ was highest in the frontal lobe and showed higher divergence in the occipital lobe (**suppl. Fig. 2B**). On an individual scale, we found that CV_i_ notably pronounced medial frontal aspects in CT-CV_i_ and showed more divergent patterns originating in the calcarine sulcus in SA-CV_i_ (suppl. **Fig. 1C**), with age-group wise distributions indicating a subtle shift towards higher dissimilarity with age (**suppl. Fig. 1C, suppl. Fig. 2C**).

To study how cortical surface metrics change in the early neonatal period, we fit a parcel-wise linear model to CT, SA, and CV_i_ metrics and determined changes with PMA, GAB, and sex across groups. Across all individuals, CT increased with PMA, most notably along the central sulcus, and age effects on SA revealed decreases along the calcarine sulcus (**Fig. 2A**). CV_i_ results, both CT-CV_i_ and SA-CV_i_, mirrored the effects of local metrics in inverse directions: CT-CV_i_ decreased with PMA in pre- and postcentral parcels, indicating increasing dissimilarity in areas of thickening CT. SA-CV_i_ changes disclosed significant increases hinting towards increasing similarity around the calcarine sulcus. We contextualized these changes using five functional neonatal networks described by Molloy and Saygin [53]. To this end, we assigned each cortical parcel to a functional network and plotted parcels that significantly changed with PMA (p_FDR_<0.05) by the network they primarily participate in. Here, we found that more unimodal networks, specifically the somatomotor and visual networks, exhibited CT-CV_i_ decreases, whereas more transmodal networks, such as the default mode network (DMN), were characterized by CT-CV_i_ increases. These changes mirrored CT effects, hinting towards an increasing dissimilarity of parcels in unimodal areas. SA-CV_i_ and SA revealed more distributed effects across functional networks (**Fig. 2B**). GAB effects mostly temporal CT increases and frontal CT decreases with concurrent CT-CV _i_ changes, with SA and SA-CV_i_ revealing more congruent decreases in medial frontal areas (**Suppl. Fig. 3**). We then assessed the influence of Apgar scores at one and five minutes (distribution within groups is shown in **Fig. 2C** and **Table 1**) on all metrics, again controlling for PMA, GAB, sex, and global metric. Here, we found statistically significant changes with Apgar scores at one minute, specifically CT-CV_i_ increases around the right precuneus, accompanied by CT increases in the right temporal lobe and precuneus. SA-CV_i_ showed a localized decrease in the right inferior temporal lobe with no associated changes in SA. The distribution of delivery modes showed the highest percentage of term-born neonates was delivered vaginally, followed by emergency caesarean section in labor, whereas preterm-born newborns had an equally high percentage of vaginal delivery and emergency caesarean section not in labor (**Fig. 2D**). When assessing the influence of birth method as compared to vaginal birth on covariance metrics, we found disseminated changes highlighting the anterior pole in CT increases and CT-CV_i_ decreases primarily along the left convexity with elective caesarean section. In addition, we found a localized significant decrease in SA with ventouse instrumental delivery in the right parietal lobe, and an SA-CV_i_ decrease in the left precuneus with ventouse instrumental delivery (**Fig. 2D**).

### Group gradients and contextualization to archetypal axes

We then sought to contextualize age-group-wise patterns of brain structural expansion. To understand how age-related changes over- and under-vary in relation to a birth-time norm, we determined the parcel-wise correlation of CV_g_ metrics to a birth time reference derived from n=89 term-born, 40-week PMA neonates. We found an inverse U-shaped pattern of correlation over age in the term group, whereas metrics in the preterm group correlated strikingly lower and at a stable level with this template (**Fig. 3A**). Parcel-wise distributions are visualized on the cortical surface in **supplemental fig. 4.** Of note, total intracranial volume values were not significantly different between groups (t=0.262, p=0.793).

**Figure 3.**
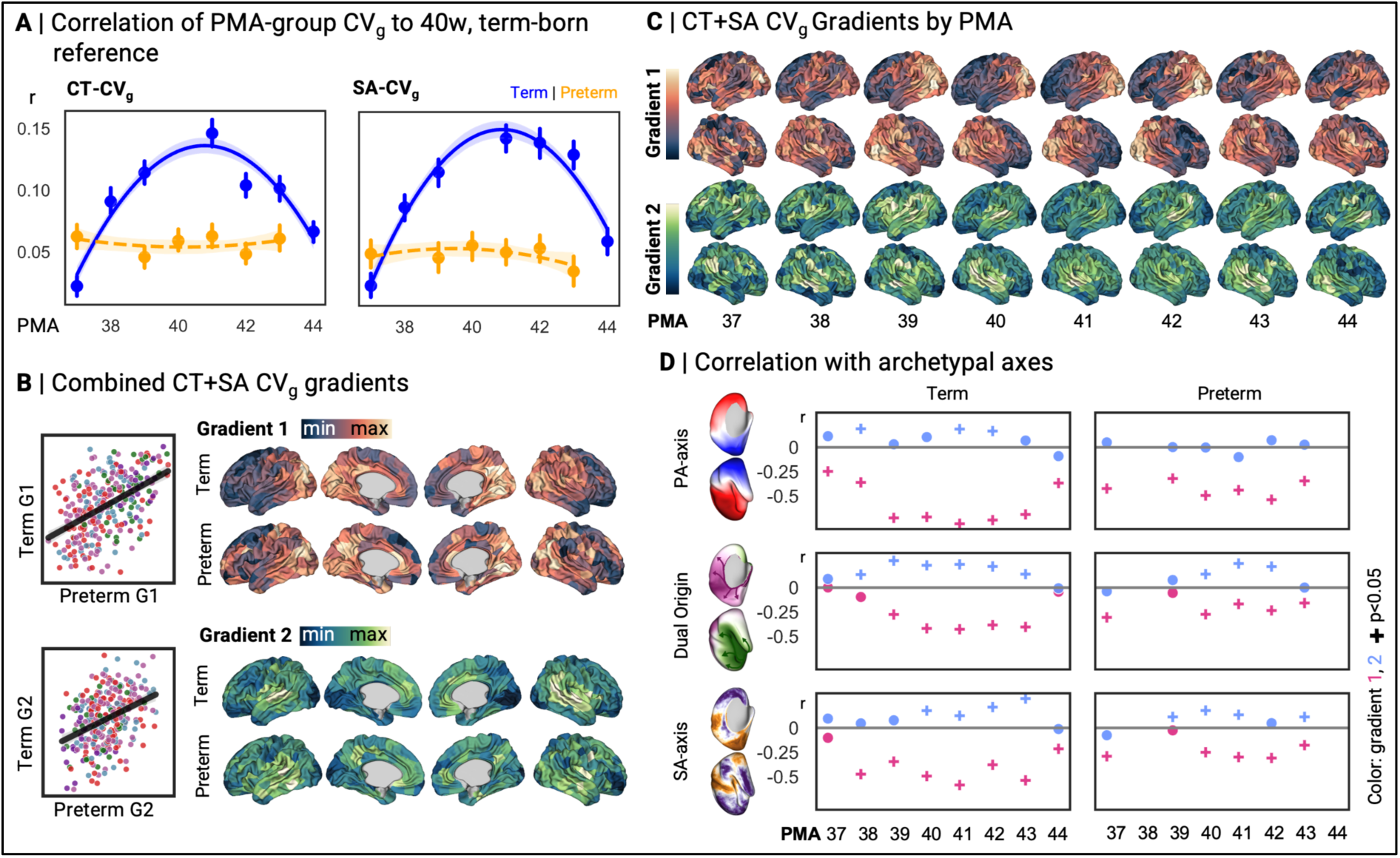
**A** Correlation of CV_g_ metrics in each age cohort to a birth time reference (derived from all 40w PMA, term-born neonates). **B** Combined CT+SA CVg gradients (joined via fusion, i.e. stacking of matrices), of term and preterm reference groups, scatterplots show correspondence with each parcel colored by their functional network (Molloy and Saygin, purple = visual, blue = somatomotor, green.= higher-order visual, pink = ventral attention network, red = default mode network, see Fig. 2 B). **C** Gradients (first and second principal component from diffusion embeddings-based representation of CT and SA matrices) in term neonates of each PMA group. **D** Correlation of gradients to archetypal brain organizational axes, from top to bottom: posterior-anterior axis, dual origin phylogenetic axis, and sensorimotor-association axis. Crosses indicate statistical significance after correcting for spatial autocorrelation (permutation-based p<0.05, i.e. observed r exceeds r in 5000 variogram-based null models).

To extract major organizational trajectories, we aggregated CT-CV_g_ and SA-CV_g_ and applied diffusion embedding for dimensionality reduction. In the resulting primary and secondary gradients, we observed an posterior-anterior pattern that appeared anchored in the occipital cortex, and secondary gradients extended from occipital areas towards lateral frontotemporal and insular regions in term-born neonates, whereas the preterm-born group disclosed more disseminated patterns with remarkably higher primary gradient scores in frontal aspects, and secondary gradients extending further into the medial parietal lobe (**Fig. 3B**). Across age, primary gradients show a shift towards a more inferior-superior distribution in the youngest and oldest groups (**Fig. 3C**).

We then tested the correlation of each age group’s gradient to archetypal brain organizational axes. Overall, we observed that gradients derived from term-born neonates exhibited higher absolute correlation coefficients with all three axes, reaching statistical significance in permutation-based testing against 5000 null models for posterior-anterior and sensorimotor-association patterns (**Fig. 3D**). Within term-born neonates, we found that correlation coefficients to any axes followed a U-shaped pattern, where the youngest and oldest age groups exhibited lower absolute correlation to the selected archetypal axes as compared to 39-42 weeks PMA groups. Preterm PMA groups of 38- and 44-weeks PMA were excluded from analysis as their sample size was insufficient. Of note, we did not observe a statistically significant correlation with dual origin patterns. In robustness analyses, we repeated correlation tests by considering a binarized dual origin territory map (**suppl. Fig. 5, suppl. Table 1**), revealing largely similar patterns in both continuous and binarized approaches.

In addition to this primary approach leveraging joint CT+SA-CV_g_ matrices, we conducted robustness analyses using CT-CV_g_ and SA-CV_g_ matrices separately to extract gradients via diffusion embedding. The resulting patterns largely mirrored posterior-anterior patterns similar to our primary joint analysis (**suppl. Fig. 6**). While overall posterior-anterior patterns in primary gradients appeared similar, SA-CV_g_ stressed the calcarine sulcus and occipital cortex, and CT-CV_g_ gradients were more smoothly distributed along posterior-anterior directions.

We then repeated this analysis using CT-CV_g_ and SA-CV_g_ gradients separately without fusion. Here, we found that CT-CV_g_ gradients showed a stringent posterior-anterior distribution, whereas primary SA-CV_g_ gradients were remarkably anchored in the occipital lobe, radiating laterally and anteriorly (**suppl. Fig. 6**). Then, we repeated correlation testing to posterior-anterior, dual origin, and sensorimotor-association axes, showing largely congruent patterns with fused analysis as term-borns showed higher correspondence to these patterns, although not reaching statistical significance in most age groups (**suppl. Fig. 7**).

### Individual-level covariance gradients

Finally, to complement group-level findings and results from surface-based linear models with whole-brain organizational representations, we computed lower-dimensional components of individual CT-CV_i_ and SA-CV_i_ profiles. To ensure common directionality, we aligned individual gradients to a group-average gradient derived from all term-born neonates at 40w PMA. The resulting primary gradients anchored in lateral frontal and parieto-temporal regions, radiating towards the medial frontal lobe; secondary gradients appeared to run from the insula to medial frontal regions. Reference gradients are shown in **Fig. 4A**. We then applied a linear model to individual gradients to test for the effects of age, GAB, and sex, revealing largely congruent increases with PMA and GAB along the left medial wall, with additional significant increases located in the frontal and temporal lobes (**Fig 4B, C**), indicating a shift of primary motor regions to the latter end of the primary gradient, along with a transition of the left temporoparietal junction to higher secondary gradient scores.

**Figure 4.**
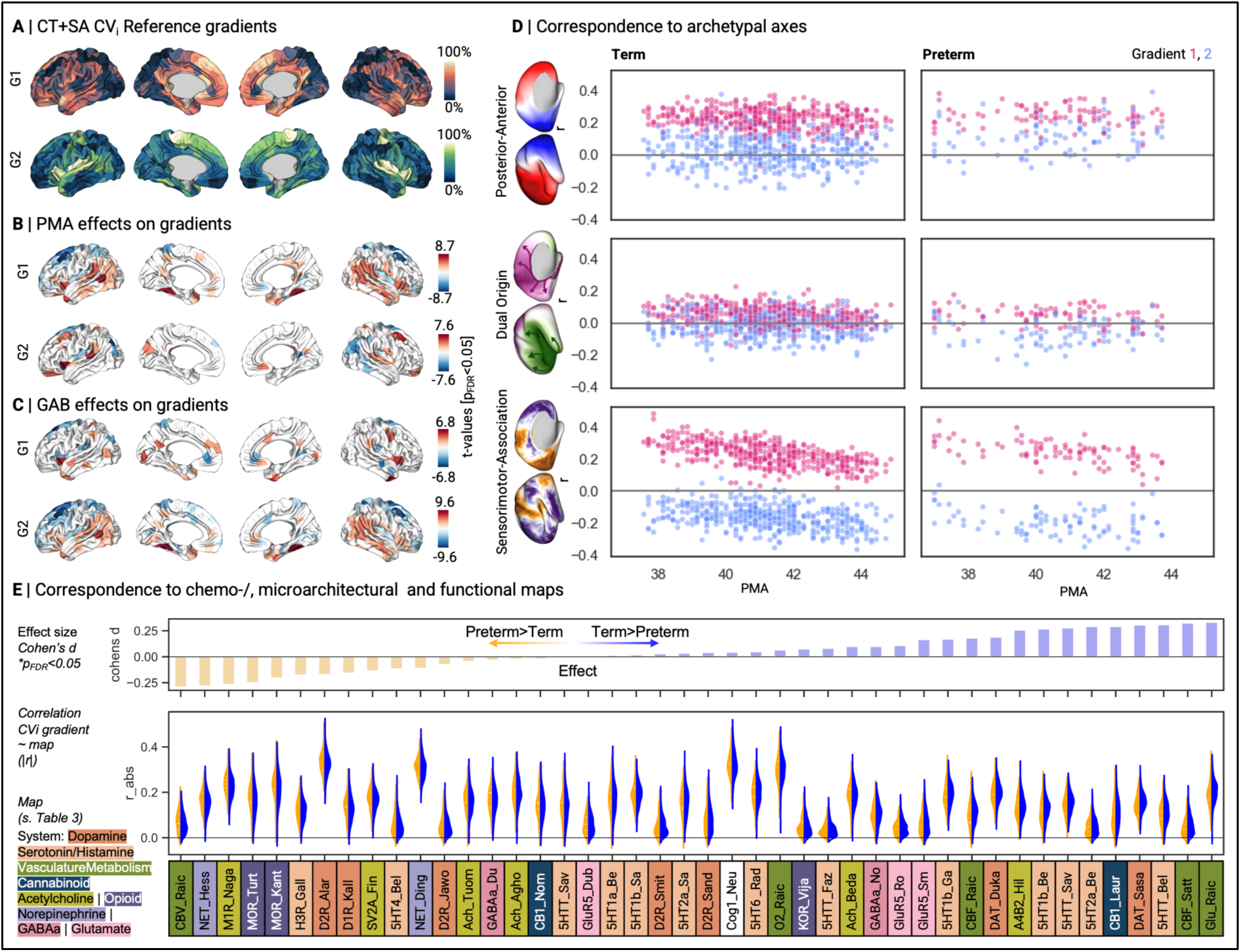
**A** Reference gradients derived from mean CVi matrices of all term-born, 40w PMA neonates, used for alignment. **B** Outcomes of linear model testing differences with postmernstrual age (PMA) and **C** gestational age at birth (GAB) in normalized gradient loadings. **D** Correspondence to selected archetypal brain organizational axes at individual-level granularity. The graph shows the correlation strength (r) of aligned individual CT+SA-CVi gradients to the posterior-anterior axis, dual origin map, and sensorimotor-association axis, respectively, with colors reflecting the correspondence of each individual’s primary (pink) and secondary gradient (blue) correlation coefficient, respectively. No significance testing was performed. **E** Correspondence to neuroarchitectural maps, assessed by testing each individual’s CT+SA-CVi map’s correlation to selected microarchitectural maps and determining an effect size estimate of r-to-z-transformed correlation coefficients between term- and preterm-born neonates. A glossary and references for maps is given in **Table 3**. Upper panel: bars marked with a star represent significant differences (p_FDR_<0.05) in correlation estimates between groups. Distribution of correlation estimates is given below, blue = term-born, yellow = preterm born. A list of maps, references and respective methods is given in table 3. CVi = individual covariance, FDR = false discovery rate, GAB = gestational age at birth, PMA = postmenstrual age

Next, we tested how these trajectories corresponded to archetypal organizational axes. Here, we found that term and preterm-born groups revealed largely similar trajectories (**Fig. 4D**). Notably, across age, correlation to the geometric axis appeared to remain stable, whereas the dual origin pattern appeared to decrease, and the correspondence of the primary gradient to the sensorimotor-association pattern revealed slightly decreasing correlation with PMA (**Fig. 4D**). We then repeated all analyses with individual CT-CV_i_ and SA-CV_i_ gradients, respectively. In linear models, we found pervasive changes in CT-CV_i_-gradients, especially along the central sulcus (**suppl. Fig. 8B, C**), whereas SA-CV_i_ changes were less pronounced (**suppl. Fig. 8F, G**). In relation to archetypal axes, CT-CV_i_ gradients correlated notably strong to dual origin patterns and showed an increasing absolute correlation strength with PMA to the sensorimotor-association axis (**suppl. Fig. 8D**). SA-CV_i_ gradients, however, showed lower correspondence to the dual origin pattern and a decreasing correspondence with the sensorimotor-association axis with PMA (**suppl. Fig. 8H**).

To contextualize neonatal gradients beyond *a priori* selected archetypal axes, we aimed to test how neonatal brain expansion corresponds to normative adult brain microstructure and chemoarchitecture patterns. To this end, we leveraged spatial patterns from *neuromaps* and determined their correspondence to each individual’s primary gradient. Then, we determined the term vs. preterm effect size of differences in r-to-z-transformed correlation coefficients, identifying those maps that correlate better to term- and preterm-born individual covariance matrices, respectively. Maps that correlated significantly higher (t-test between correlation coefficients, p_FDR_<0.05) with CV_i_ gradients of preterm newborns as compared to term neonates were linked to cerebral blood volume, a norepinephrine- and acetyclcholine transporter, whereas patterns corresponding more to term-born patterns encompassed cerebral blood flow and glucose metabolism maps (**Fig. 4E**).

### Robustness analyses

For both group-level and individual gradients, we conducted robustness analyses to test whether linear dimensionality reduction using Laplacian eigenmaps (LE) results in similar gradient patterns. For each reference gradient derived from all term-born neonates at 40-week PMA, we repeated gradient decomposition using LE and determined the correlation of the resulting components to nonlinear diffusion embedding-derived gradients from our primary analysis. Both for CV_g_ and CV_i_, the resulting primary and secondary gradients were strongly correlated (**supplemental table 1,** Pearson’s r>0.9 across all source metrics). Moreover, we tested gradient stability across different sparsity thresholds (no sparsity in main approach; thresholds 0.1 to 0.9 in 0.1 inclinations in robustness testing). Pearson’s correlation coefficients to the main approach were considerably high across all thresholds (**supplemental table 2**). All major analyses based on joint CT+SA-CV metrics were repeated in each covariance metric in isolation and are attached in the supplements, with specific references given with each analysis.

## DISCUSSION

The brain’s relative immaturity at birth presents a conditional benefit: while increasing vulnerability to early stressors [6], it also allows unique adaptability to changing environmental demands throughout development [5], [23]. Describing brain structural maturation in the early neonatal period is crucial to understanding disrupted consolidation of organizational axes throughout neurodevelopment [96], [97]. We here complemented morphological features from structural MRI of neonates around term-equivalent age with covariance metrics mirroring interregional differentiation, to investigate the synchrony of regional brain maturation in the neonatal period. Across PMA and GAB, we observed that structural changes followed functional patterns: significant changes, especially in CT-CV_i_ and local CT exhibited opposite directional trends in unimodal versus transmodal systems. Likewise, individual-level gradients, i.e., representations of CT-CV_i_ and SA-CV_i_ covariance profiles extracted using dimensionality reduction, showed an increasing correlation with a sensorimotor-association axis with PMA. Across analyses, our results underline a progressive alignment of structural marker-derived interregional synchrony with functionally informed patterns in early development.

We here leveraged two different measures of co-maturation that reflect individual and group-level synchrony to understand how both individual-level and age group-wise patterns develop in the neonatal period. First, we investigated CV_i_, a normalized similarity function first introduced by Wee et al. [38]. We here interpret CV_i_ increases as higher similarity between parcels and covariance decreases with age as an increasing differentiation between cortical parcels. The original description of CV_i_ leveraged CT in older adults [38], impeding the comparison of our results to age-appropriate literature; however, the spatial congruence with raw metrics strengthens the plausibility of CV_i_ patterns even in the absence of age-specific references. Vis-à-vis individual covariance, we utilized group-level CV_g_ to reflect larger-scale, population-level trends in cortical co-maturation. Across covariance metrics, we found that intrinsic trajectories were largely anchored in the occipital cortex, radiating anteriorly. As such, both covariance approaches used here appear to converge in largely similar patterns, further underlining the biological plausibility of our results. Overall, we observed that group-level covariance gradients appeared to correspond to the posterior-anterior and sensorimotor-association axis. In individual covariance gradients, we found the association to the posterior-anterior distribution to remain stable across age, whereas correspondence to a sensorimotor-association axis disclosed an increase with PMA. In group-wise analysis, this effect appeared to diminish in oldest age groups, possibly mediated by smaller sample sizes. These patterns proved consistent in confirmatory analyses using non-fused CT- and SA-CV gradients, and by testing the reliability of gradient decomposition by showing a high congruence of gradients from linear and nonlinear dimensionality reduction. Taken together, these results hint towards both patterns as primary spatial trajectories for early life brain structural expansion, yet little allo-cortical differentiation. Overall, these changes potentially reflect increasing cortical specialization and differential growth, underscoring regions that are particularly active in response to external stimuli [28], [53], [98].

Brain structure and function are closely entangled, i.e., local variability in cortical microarchitecture covaries with whole-brain functional patterns [23], [39], [40]. At birth, synaptogenesis increases sharply, followed by an increase in overall connectivity and neuronal interaction [14], [16], [99]. After birth, volume expansion continues until adulthood, mostly driven by white matter development [14], [100]. Cellular mechanisms underlying cortical expansion during late gestation in the neonatal period, i.e., the “big bang” of exuberant connection formation and synaptogenesis [14], [99], build a foundation for future whole-brain connectivity patterns and efficient signal processing through functional hierarchies. Local CT and SA increases are hypothesized to stem from cellular proliferation and connection formation in early development [14], [101], [102]; as such, the observed changes in CV_i_ likely reflect spatial differentiation in a neonatal cortex that adapts to the extrauterine environment and novel stimuli [3], [48], while the underlying genetic and biomolecular underpinnings remain opaque in the current, structural imaging-based approach.

Throughout neurodevelopment, brain areas undergo functional specialization, allowing for hierarchical segregation and integration of information processing [21], [27], [28], [30]. Thalamocortical connectivity, handling information in- and output sensory organs [103], increases in the perinatal period [16], [21], [104], [105] and remains a mainstay of primary signal processing throughout life [106], [107]. Previous studies leveraging functional MRI in neonates have demonstrated primary gradients extending from pericentral sensorimotor to visual areas [21], notably differentiating primary functional regions and mirroring a secondary functional gradient known from previous research in healthy adults [55]. Secondary functional gradients in neonates paralleled a cortical thickness-like posterior-anterior distribution [23], [108], potentially reflecting the development of transmodal areas, and their functional segregation from primary input [20], [27]. These patterns have been hypothesized to align with spatial patterns of thalamic connections, coinciding with an increase in thalamocortical connectivity in the perinatal period [17], [105]. Throughout the lifespan, the thalamus remains a central contributor to connectivity maturation patterns along the sensorimotor-association axis, mirroring functional refinement and development along this trajectory throughout childhood and adolescence [1], [28], [106], [107]. While immature, functional networks active at rest that resemble the functional organization of the adult neocortex have been described in neonates [53], [109]. We found that these regions, specifically in the visual and sensorimotor cortex, displayed structural expansion and differentiation at term-equivalent age, in alignment with increased signal processing along thalamocortical connections.

In addition to functional pattern associations, we found a significant correlation with a posterior-anterior axis throughout age groups. Of note, given the observed accentuation of primary sensory areas, including visual systems in the occipital cortex, this correlation might be partly mediated by functional patterns. The concepts of posterior-anterior and sensorimotor-association patterns have been shown to intersect, stemming from complementary genetic programs [35]. Moreover, geometry has been proposed as a central fundamental constraint of human brain function [110], [111], [112], [113], with the posterior-anterior axis constructed here paralleling the second eigenmode described by Pang et al. [110]. Notably, we did not observe strong significant correlations to a phylogenetically informed dual origin pattern, whereas significant correlations to sensorimotor pattern further support thalamocortical connectivity shifts as an important driver of cortical co-maturation in the early neonatal period.

Of note, while we found a persistent association to posterior-anterior and sensorimotor-association axes in group-level CV_g_ gradients, this correspondence was higher and more consistent across age in term-as compared to preterm-born neonates. Overall, preterm-borns revealed lower correlation to a birth-term norm and to archetypal brain organizational trajectories, underlining fundamental differences in structural expansion of preterm neonates that appear more disconnected from geometry or function. Taken together, these findings underline different developmental trajectories in preterm-born neonates that are not simply temporally shifted, but inherently different from structural expansion in term-born infants. Our observations align with previous reports of pervasive brain changes and altered developmental trajectories after preterm birth [8], [97], [114]. However, given considerably lower sample sizes in preterm-born groups, this effect might be mediated by sample size and warrants further exploration.

To further elucidate differences in maturation patterns between term- and preterm-born groups, we assessed how their primary gradients correlated to chemoarchitectural and metabolic maps of the brain, using individual gradients to overcome sample size limitations in the computation of group matrices. Here, we found there was a higher number of maps that correlated stronger with individual covariance gradients from term infants relative to pre-term infants. Among the patterns that were more strongly associated with term-borns co-maturation patterns were cerebrovascular and metabolic maps, with notably different effects in cerebral blood flow and volume to brain expansion of term- and preterm-born infants respectively. These changes potentially reflect a closer adherence to cerebrovascular and microstructurally programmed trajectories in term-borns, whereas infants born prematurely show less stringent structural expansion that appears more detached from whole-brain, adult organizational patterns. While this computational, exploratory approach does not explain the underlying physiology behind differential expansion with birth timing on a cellular level, it hints towards the involvement of brain metabolic pathways in the structural expansion after birth. Of note, as pediatric reference maps are not readily available to date, chemoarchitectural maps used in this study stem from adult populations and thus do not reflect the actual microarchitecture in neonatal brains; rather, they indicate spatial patterns that are to emerge through continued neurodevelopment. However, given that the spatial distribution of receptor patterns is co-determined by genetic programming [77], [115], early growth disruptions convergent with receptor expression patterns might still hint towards long-term atypicality within these organizational trajectories.

### Strengths and limitations

Through the use of an existing open neuroimaging dataset, we were able to include a large number of newborns at different ages, maximizing the generalizability of our results. As we used a sample scanned at and around term-equivalent age, preterm-born neonates were chronologically older than term-borns, introducing collinearity and impeding the comparability of term- and preterm-born groups. Where possible, we focused analyses on term-born neonates as a reference, describing qualitative differences in developmental trajectories in preterm-born infants.

While our results provide a structural imaging substrate of early aberrant co-maturation at term-equivalent age, the functional correlates of these changes remain unknown. In addition, results here focus primarily on large-scale brain organizational trajectories in term-born neonates, while the heterogeneous preterm population is aggregated to show general deviations. However, while we here regard all infants born before 37 weeks of gestational age as preterm; the variety of mitigating factors in premature development warrant future investigation of preterm-born infants in further detail, including health and environmental factors.

## CONCLUSION

We here utilized covariance measures to capture interregional synchrony of structural cortical organization in the neonatal period. Our results highlight profound morphological changes that link to functional and geometrical axes of brain organization. We find a robust and increasing correspondence of intrinsic co-maturation patterns with an archetypal sensorimotor-association axis, hypothesized to reflect developing functional patterns in alignment with previous research [20], [97], [100]. In addition, we show that preterm-born infants exhibit inherently different patterns of structural expansion that appear to be largely independent from archetypal organizational patterns. We confirm these results from combined CT and SA covariance metrics on group- and individual scales in CT and SA in isolation, showing largely convergent effects. Future directions include the contextualization of our findings to functional imaging and clinical outcome ratings.

## Acknowledgments

Data were obtained from the developing Human Connectome Project (https://www.developingconnectome.org/), which is openly available in the NIMH data archive (https://nda.nih.gov/edit_collection.html?id=3955).

## Abbreviations

CT: cortical thickness
CV: covariance
CV_g_: group-level covariance
CV_i_: individual covariance
FDR: false discovery rate
GAB: gestational age at birth
ICV: intracranial volume
MRI: magnetic resonance imaging
PET: positron emission tomography
PMA: postmenstrual age
SA: surface area,
SD: standard deviation

## Code availability

github.com/claraweber/babybrains_structexp

