## Supplemental Material for "Structural Brain Co-Maturation in the Neonatal Period"

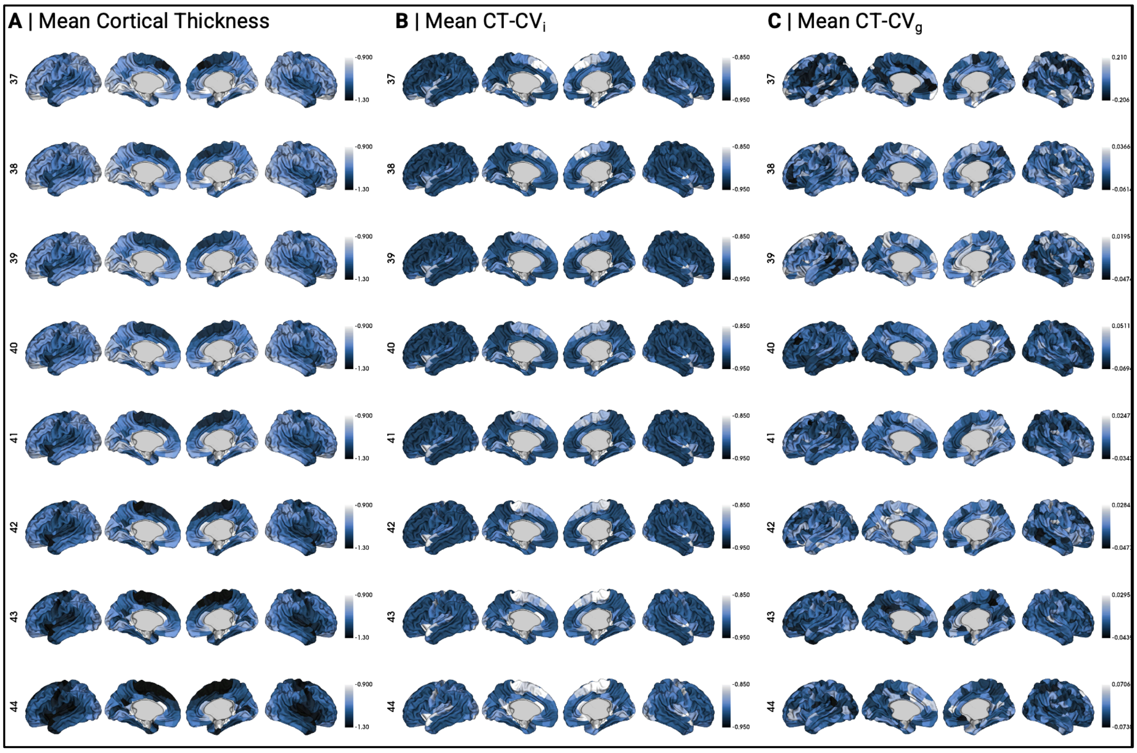


**Suppl. Figure 1. A** Mean CT, **B** CT-CV$_i$ and **C** mean CT-CV$_g$ in each PMA cohort. CT = cortical thickness, CT-CV$_g$ = group-level covariance based on CT, CT-CV$_i$ = individual covariance measure based on CT, PMA = postmenstrual age.

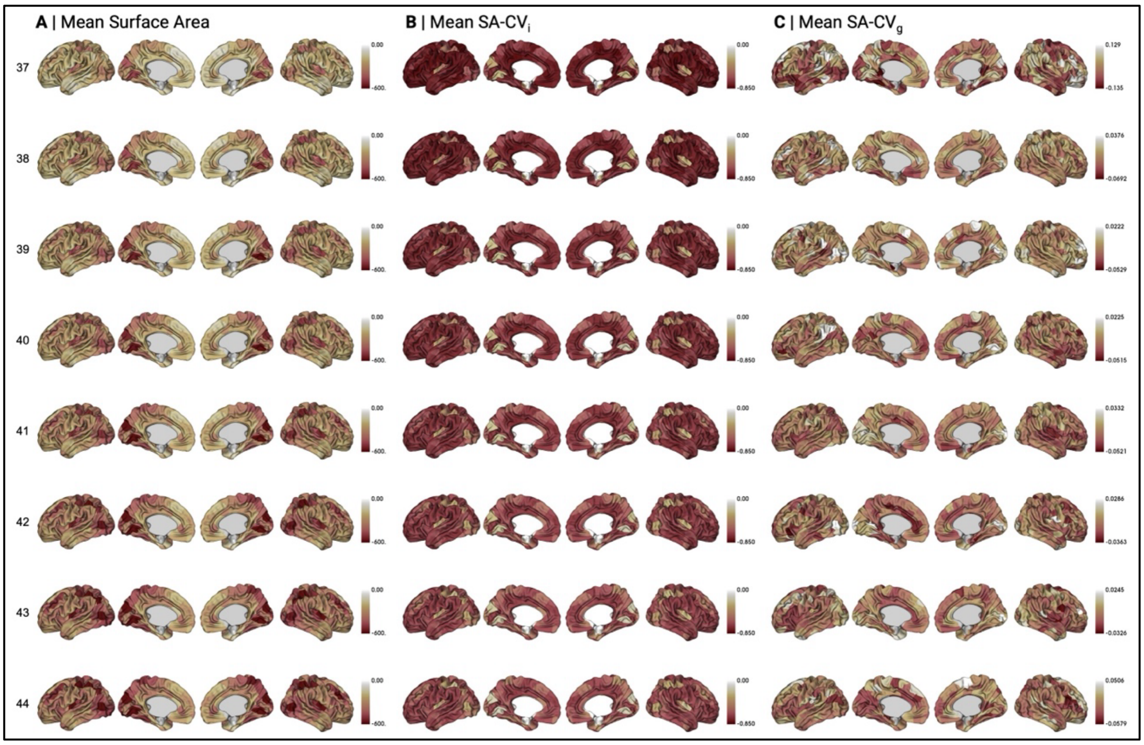

**SUPPL. FIGURE 2. A** Mean SA, **B** SA-$CV_i$ and **C** mean SA-$CV_g$ in each PMA cohort. SA = surface area, SA-$CV_g$ = group-level covariance based on SA, SA-$CV_i$ = individual covariance measure based on SA, PMA = postmenstrual age.

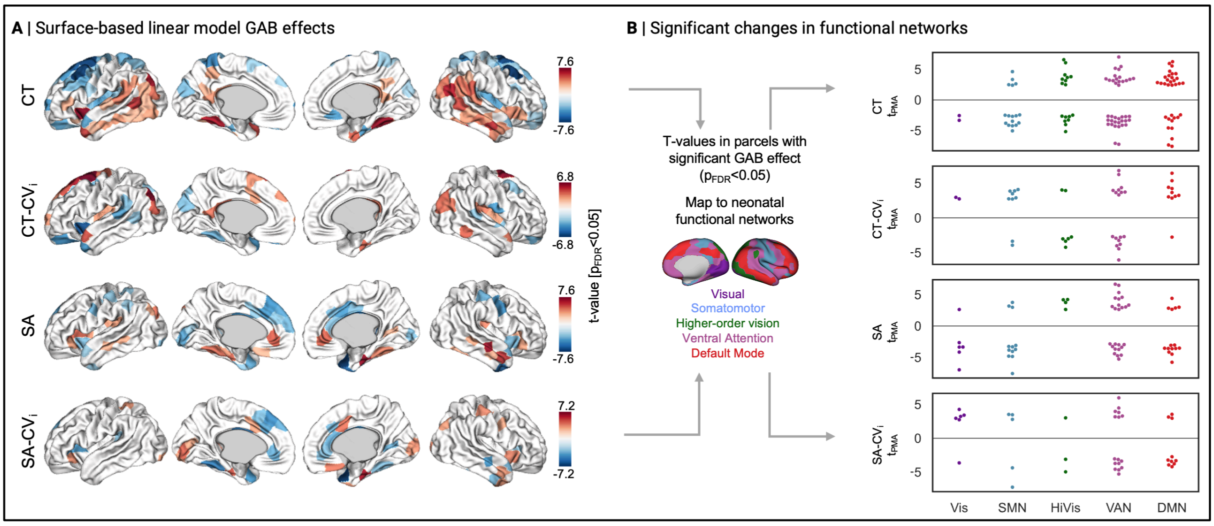


**SUPPL. FIGURE 3. A** Effect of GAB in **B** neonatal functional networks, parallel to Fig. 2. SA = surface area, SA-$CV_g$ = group-level covariance based on SA, SA-$CV_i$ = individual covariance measure based on SA, GAB = gestational age at birth.

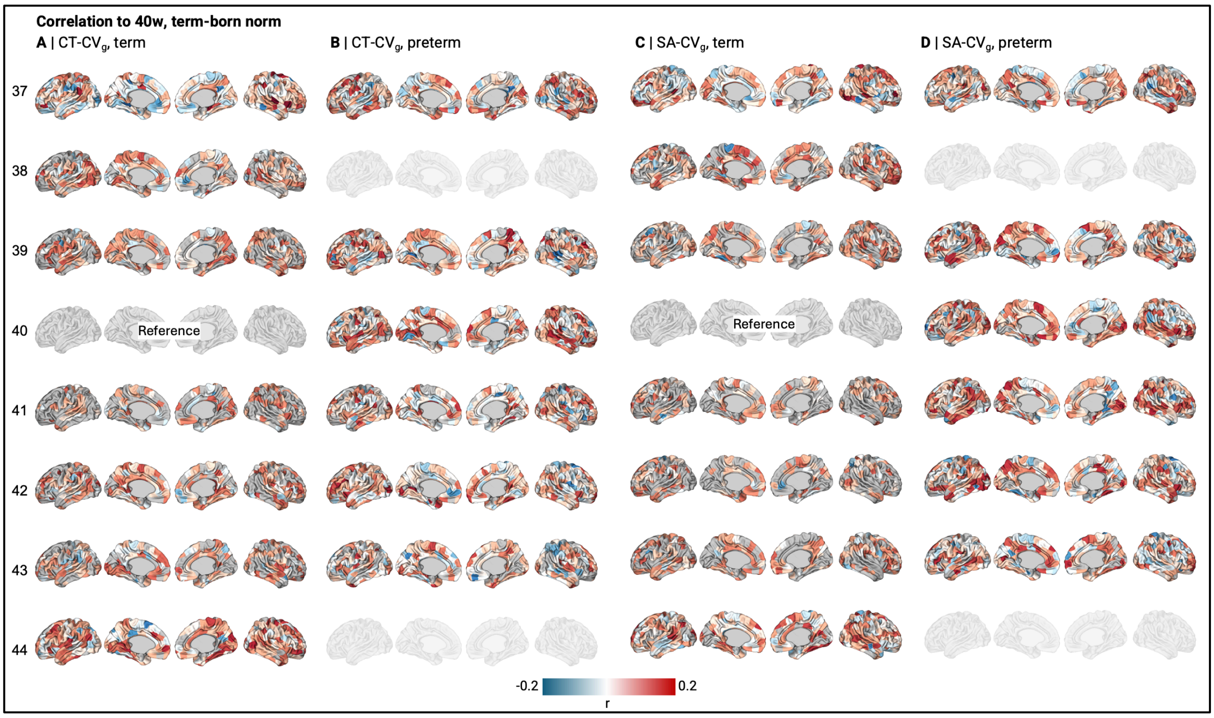


**SUPPL. FIGURE 4.** Correlation of $CV_g$ metrics to 40w birth time norm in CT-$CV_g$ and SA-$CV_g$ separately, and stratified by term status. $p<0.05$ is marked as grey. 40w term as the reference group is excluded, so are 38w and 44w preterm groups, as not enough subjects were available in these age groups

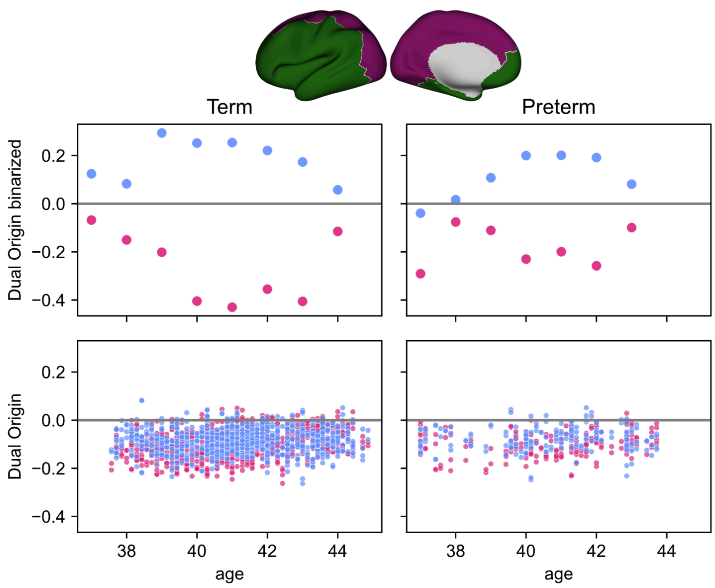


**SUPPL. FIGURE 5.** Robustness analysis for dual origin pattern correlation. Graphic shows Spearman correlation of primary (pink) and secondary (blue) gradients to binarized dual origin pattern for group (upper panel) and individual (lower panel) covariance. In group covariance, statistical significance was corrected for spatial autocorrelation in 5,000 variogram-based permutations. Scatter shape indicates significance (· = n.s., × = $p_{perm}$<0.05).

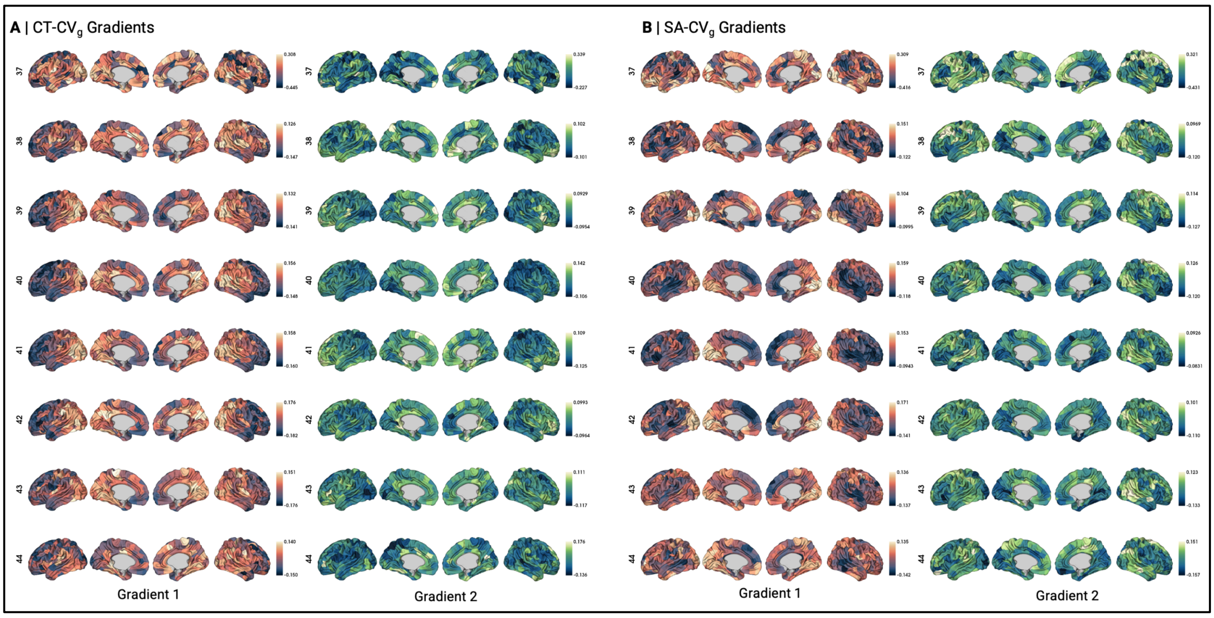


**SUPPL. FIGURE 6.** Primary and secondary gradients derived from CT-CVg and SA-CVg by PMA group.

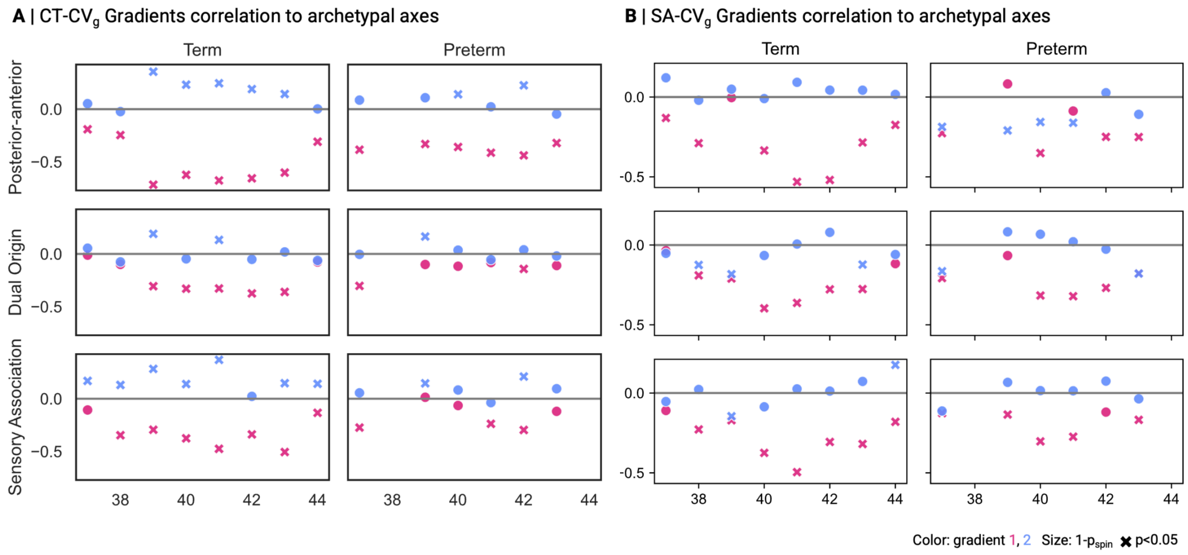


**SUPPL. FIGURE 7.** Correlation of CT- (**A**) and SA-CVg (**B**) primary (pink) and secondary (blue) gradients to three archetypal brain organizational axes. Statistical significance after correction for spatial autocorrelation is shown by point shape.

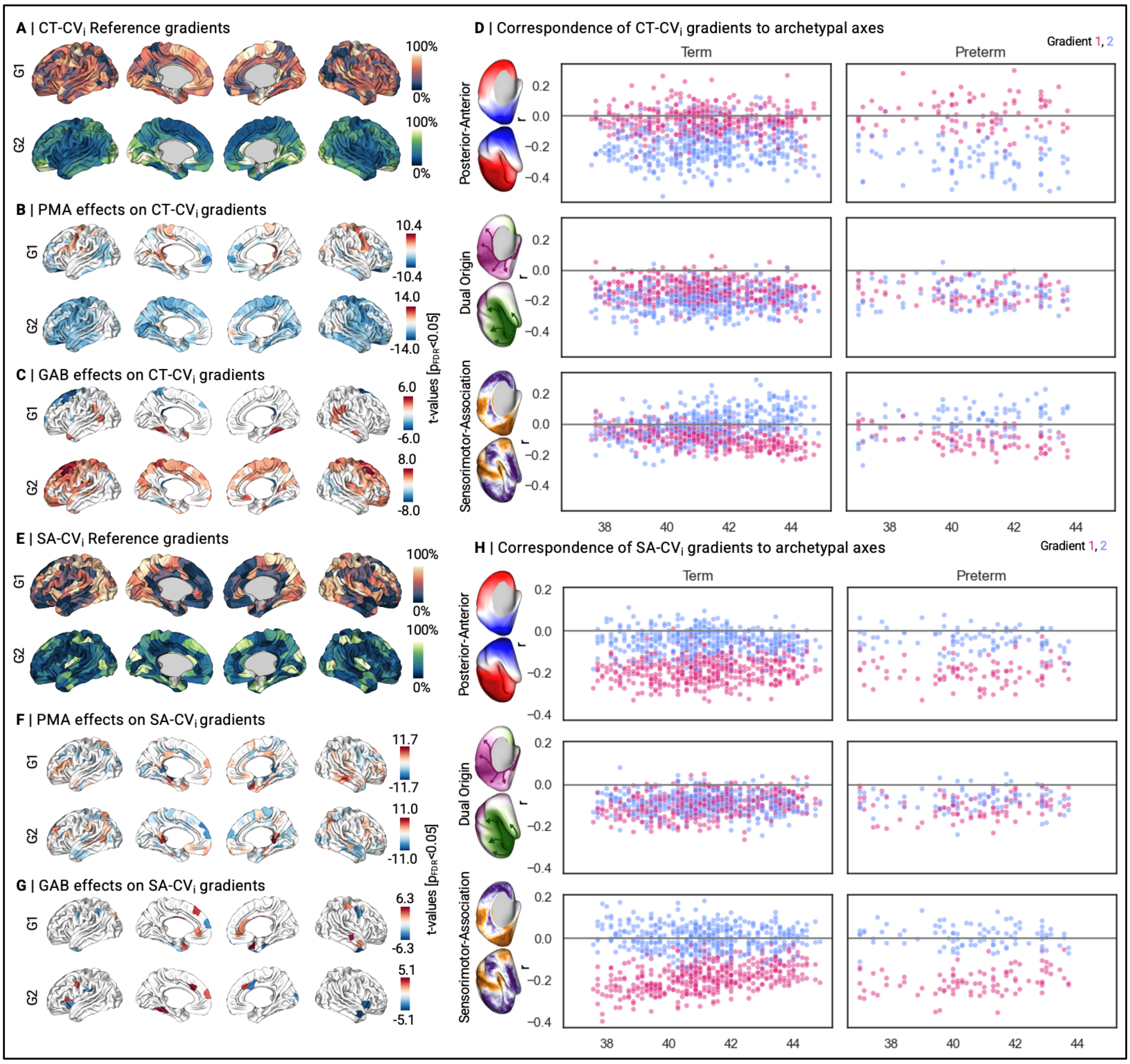

**SUPPL. FIGURE 8.** Individual gradients from CT (upper panels) and SA (lower panels). **A/E** Reference gradients derived from an averaged $CV_i$ matrix of all 40w, term-born neonates. These gradients were used for alignment of individual gradients. **B/C** and **F/G** show significant ($p_{FDR}$<0.05) changes (t-values) in normalized, aligned gradients with postmenstrual age (PMA) and gestational age at birth (GAB), correcting for sex. **D/H** Correlation of each individual first (pink) and second (blue) gradient to three archetypal brain organizational axes.

**SUPPL. TABLE 1.** Pearson correlation of gradients derived by diffusion embedding (main approach) versus Laplacian eigenmapping.

| | **CT+SA CVi** | | **CT+SA CVg** | |
|---|---|---|---|---|
| **Gradient** | **r** | **p** | **r** | **p** |
| 1 | >0.999 | <0.001 | >0.999 | <0.001 |
| 2 | 0.991 | <0.001 | >0.999 | <0.001 |
| | **CT CVi** | | **CT CVg** | |
| **Gradient** | **r** | **p** | **r** | **p** |
| 1 | 0.992 | <0.001 | >0.999 | <0.001 |
| 2 | 0.959 | <0.001 | >0.999 | <0.001 |
| | **SA CVi** | | **SA CVg** | |
| **Gradient** | **r** | **p** | **r** | **p** |
| 1 | >0.999 | <0.001 | >0.999 | <0.001 |
| 2 | >0.999 | <0.001 | >0.999 | <0.001 |

**SUPPL. TABLE 2.** Pearson correlation of gradients at different sparsity thresholds to the main approach, where no sparsity was applied.

| | | **Correlation to CT+SA-$CV_i$** | | **Correlation to CT+SA-$CV_g$** | |
|---|---|---|---|---|---|
| **Sparsity** | **Gradient** | **r** | **p** | **r** | **p** |
| 0 | Main approach/reference | | | | |
| 0.1 | 1 | 0.999 | <0.001 | >0.999 | <0.001 |
| | 2 | 0.985 | <0.001 | 0.998 | <0.001 |
| 0.2 | 1 | 0.994 | <0.001 | >0.999 | <0.001 |
| | 2 | 0.945 | <0.001 | 0.998 | <0.001 |
| 0.3 | 1 | 0.990 | <0.001 | 0.999 | <0.001 |
| | 2 | 0.912 | <0.001 | 0.998 | <0.001 |
| 0.4 | 1 | -0.988 | <0.001 | 0.997 | <0.001 |
| | 2 | 0.913 | <0.001 | 0.990 | <0.001 |
| 0.5 | 1 | 0.976 | <0.001 | 0.992 | <0.001 |
| | 2 | 0.912 | <0.001 | 0.978 | <0.001 |
| 0.6 | 1 | 0.940 | <0.001 | 0.985 | <0.001 |
| | 2 | 0.871 | <0.001 | 0.964 | <0.001 |
| 0.7 | 1 | 0.901 | <0.001 | 0.973 | <0.001 |
| | 2 | 0.814 | <0.001 | 0.903 | <0.001 |
| 0.8 | 1 | 0.858 | <0.001 | 0.946 | <0.001 |
| | 2 | 0.720 | <0.001 | 0.910 | <0.001 |
| 0.9 | 1 | 0.796 | <0.001 | 0.910 | <0.001 |
| | 2 | -0.441 | <0.001 | 0.890 | <0.001 |
